# Translocator protein is a marker of activated microglia in rodent models but not human neurodegenerative diseases

**DOI:** 10.1101/2022.05.11.491453

**Authors:** Erik Nutma, Nurun Fancy, Maria Weinert, Manuel C. Marzin, Stergios Tsartsalis, Robert C.J. Muirhead, Irene Falk, Joy de Bruin, David Hollaus, Robin Pieterman, Jasper Anink, David Story, Siddharthan Chandran, Jiabin Tang, Maria C. Trolese, Takashi Saito, Takaomi C. Saido, Katie Wiltshire, Paula Beltran-Lobo, Alexandra Philips, Jack Antel, Luke Healy, Craig S. Moore, Caterina Bendotti, Eleonora Aronica, Carola I. Radulescu, Samuel J. Barnes, David W. Hampton, Paul van der Valk, Steven Jacobson, Paul M. Matthews, Sandra Amor, David R. Owen

**Affiliations:** Department of Pathology, Amsterdam UMC – Location VUmc, Amsterdam, the Netherlands; Department of Brain Sciences, Imperial College London, UK; UK Dementia Research Institute at Imperial College London, UK; Department of Psychiatry, University of Geneva, Switzerland; Viral Immunology Section, NIH, Bethesda, Maryland, USA; Flow and Imaging Cytometry Core Facility, NIH, Bethesda, Maryland, USA; Department of Pathology, Amsterdam UMC – Location AMC, Amsterdam, the Netherlands; Centre for Clinical Brain Sciences, The University of Edinburgh, Edinburgh, United Kingdom; Department of Neuroscience, Mario Negri Institute for Pharmacological Research IRCCS, Milan, Italy; Laboratory for Proteolytic Neuroscience, RIKEN Brain Science Institute, Wako-shi, Saitama, Japan; Department of Neurocognitive Science, Institute of Brain Science, Nagoya City University, Japan; Department of Basic and Clinical Neuroscience, Institute of Psychiatry, King’s College, London, UK; Montreal Neurological Institute, McGill University, Montreal, Canada; Division of Biomedical Sciences, Memorial University of Newfoundland, Canada; Department of Neuroscience and Trauma, Blizard Institute, Barts and the London School of Medicine & Dentistry, Queen Mary University of London, London, UK

**Keywords:** ALS, AD, MS, TSPO, microglia

## Abstract

Microglial activation plays central roles in neuro-inflammatory and neurodegenerative diseases. Positron emission tomography (PET) targeting 18kDa Translocator Protein (TSPO) is widely used for localising inflammation *in vivo*, but its quantitative interpretation remains uncertain. We show that TSPO expression increases in activated microglia in mouse brain disease models but does not change in a non-human primate disease model or in common neurodegenerative and neuroinflammatory human diseases. We describe genetic divergence in the TSPO gene promoter, consistent with the hypothesis that the increase in TSPO expression in activated myeloid cells is unique to a subset of species within the *Muroidea* superfamily of rodents. We show that TSPO is mechanistically linked to classical pro-inflammatory myeloid cell function in rodents but not humans. These data emphasise that TSPO expression in human myeloid cells is related to different phenomena than in mice, and that TSPO PET reflects density of inflammatory cells rather than activation state.

## Introduction

Neuronal-microglial signalling limits microglial inflammatory responses under homeostatic conditions^1^. The loss of this cross talk in central nervous system (CNS) pathology partly explains why microglia adopt an activated phenotype in many neurodegenerative diseases^2, 3^. Genomic, *ex vivo* and preclinical data imply that microglial activation also may contribute to neurodegeneration^4^, for example, by releasing inflammatory molecules in response to infectious or damage-related triggers^5^. These lead to both neuronal injury and, more directly, pathological phagocytosis of synapses^5, 6^. Development of tools which can reliably detect and quantify microglial activation in the living human brain has been an important goal. By enabling improved stratification and providing early pharmacodynamic readouts, these would accelerate experimental medicine studies probing disease mechanisms and early therapeutics.

Detection of 18kDa Translocator Protein (TSPO) with positron emission tomography (PET) has been widely used to quantify microglial activation *in vivo*^7^. In the last 5 years alone, there have been ∼300 clinical studies using TSPO PET to quantify microglial responses in the human brain, making it the most commonly used research imaging technique for this purpose.

The TSPO signal is not specific to microglia, and the contribution from other cell types (particularly astrocytes and endothelial cells) is increasingly acknowledged^8^. The justification for quantifying TSPO as a marker of microglial activation is based on the assumption that when microglia become activated, they adopt a classical pro-inflammatory phenotype and TSPO expression is substantially increased, ^7, 9^ ^10^. This has been demonstrated repeatedly in mice, both *in vitro* and *in vivo*^11–14^. We have shown, however, that classical proinflammatory stimulation of human microglia and macrophages *in vitro* with the TLR4 ligand lipopolysaccharide (LPS) does not induce expression of TSPO^15^. Furthermore, in multiple sclerosis (MS), TSPO does not appear to be increased in microglia with activated morphology ^16^. These data appear inconsistent with the assumption that TSPO is a marker of activated microglia in humans.

To address this issue, we performed a meta-analysis of publicly available expression array data and found that across a range of pro-inflammatory activation stimuli, TSPO expression is consistently and substantially increased in mouse, but not human macrophages and microglia *in vitro*. We then performed a comparative analysis of the TSPO promoter region in a range of mammalian species and found that the binding site for AP1 (a transcription factor which regulates macrophage activation in rodents^17^) is present in and unique to a subset of species within the *Muroidea* superfamily of rodents. Consistent with the hypothesis that this binding site is required for the increase in TSPO expression that accompanies pro-inflammatory stimulation, we show that TSPO is inducible by LPS in the rat (another *Muroidea* species with the AP1 binding site in the TSPO core promoter) but not in other mammals. Because neuronal interactions modulate microglial phenotype, we then compared microglial TSPO expression in neurodegenerative diseases affecting the brain and spinal cord (Alzheimer’s Disease (AD) and amyotrophic lateral sclerosis (ALS), respectively) as well as the classical neuroinflammatory brain disease MS which features highly activated microglia. We compared each human disease to its respective commonly used mouse models (amyloid precursor protein (*App^NL-G-F^*)^18^, tau (Tau^P301S^)^19^, superoxide dismutase 1 (SOD1^G93A^)^20^, and experimental autoimmune encephalomyelitis (EAE) in young and aged animals^21^. We also studied TSPO expression with EAE in the marmoset in conjunction with frequent MRI scanning that allowed for identification of the acute lesions which contain pro-inflammatory microglia. Consistent with the *in vitro* data, we show that in AD, ALS and MS, and in marmoset EAE, TSPO protein expression does not increase in CNS myeloid cells that express a pro-inflammatory phenotype, while expression is markedly increased in activated myeloid cells in all mouse models of these diseases. With exploration of the relative expression of TSPO in publicly available CNS single cell RNA sequencing (scRNAseq) data from brains of the human diseases and rodent models, we again show an increase in microglial TSPO gene expression in mice with proinflammatory stimuli, but not humans. Finally, using functional studies and examination of transcriptomic co-expression networks, we find that TSPO is mechanistically linked to classical pro-inflammatory myeloid cell function in rodents but not humans.

These data suggest that the commonly held assumption that TSPO PET is sensitive to microglial *activation* is true only for a subset of species within the *Muroidea* superfamily of rodents. In contrast, in humans and other mammals, it simply reflects the local density of inflammatory cells irrespective of the disease context. The clinical interpretation of the TSPO PET signal therefore needs to be revised.

## Results

### *TSPO* expression and epigenetic regulation in primary macrophages

To investigate *TSPO* gene expression changes in human and mouse a meta-analysis was performed using publicly available macrophage and microglia transcriptomic datasets upon pro-inflammatory stimulation (Fig. 1). We found 10 datasets (Fig. 1a) derived from mouse macrophages and microglia in samples from 68 mice and with inflammatory stimuli including activation with LPS, Type 1 interferon (IFN), IFNγ, and LPS plus IFNγ. We performed a meta-analysis and found that *Tspo* was upregulated under pro-inflammatory conditions (Fig. 1a). In the individual datasets, *Tspo* was significantly upregulated in 9 of the 10 experiments. We then interrogated 42 datasets from primary human macrophages and microglia involving samples from 312 participants, with stimuli including inflammatory activation with LPS, IFNγ, IL1, IL6, PolyIC, viruses, and bacteria (Fig. 1b). In the meta-analysis, there was a non-significant trend towards a *reduction* in human *TSPO* expression under pro-inflammatory conditions (Fig. 1b). In the individual datasets, *TSPO* was unchanged in 33/42 (79%) of the datasets, significantly downregulated in 8/42 (19%) and significantly upregulated in 1/42 (2%). In contrast to the findings in mice, our analysis thus suggests that TSPO expression is not upregulated in human microglia and macrophages after pro-inflammatory stimulation *in vitro*.

**Figure 1.**
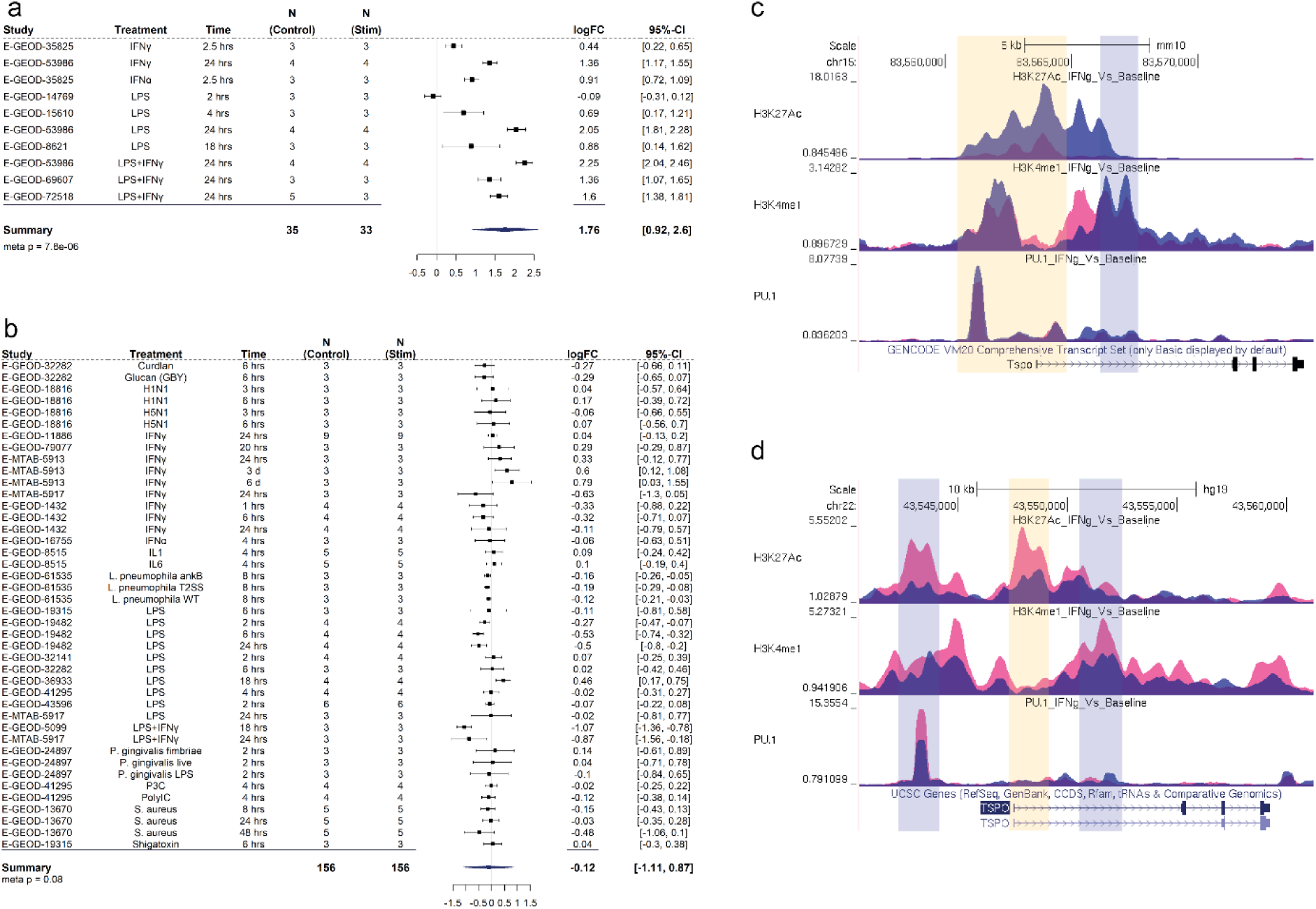
*TSPO* gene expression and epigenetic profile in human and mouse macrophages. **a,b** Forest plot of the meta-analysis for TSPO expression in **a** mouse and **b** human myeloid cells treated with a pro-inflammatory stimulus. The random-effect model was applied when combining the gene expression. The black squares represent the logFC value of each dataset. The horizontal lines indicate the 95% confidence intervals of each study. The diamond represents the pooled logFC. **c,d** ChIP-seq data, generated from **c** mouse and **d** human myeloid cells treated with IFNγ, visualisation of histone modification peaks (H3K27Ac, H3K4me3, H3K4me1) and PU.1 binding peaks at TSPO loci in IFNγ-treated (pink) and baseline (blue) conditions. Yellow vertical shading corresponds to the TSS along with promoter and light blue shading corresponds to the enhancer region of the loci.

To test whether TSPO gene expression changes are regulated at an epigenetic level, we analysed publicly available ChIP-seq datasets for histone modification in mouse and human macrophages before and after treatment with IFNγ^22^^23^ (Fig. 1c-f). Levels of H3K27Ac and H3K4me1 histone marks in the enhancer regions are associated with increased gene expression^22, 24^. While both histone modifications were increased after IFNγ treatment in TSPO promoter regions in macrophages from mouse, they were decreased in humans (Fig. 1c,d). Consistent with this epigenetic regulation, *Tspo* gene expression was upregulated in mouse macrophages after IFNγ but not in human macrophages in RNAseq data from the same set of samples (Fig. S1a).

The PU.1 transcription factor is a master regulator of macrophage proliferation and macrophage differentiation^25, 26^. Because PU.1 increases *Tspo* gene expression in the immortalised C57/BL6 mouse microglia BV-2 cell line^27^, we next investigated whether *TSPO* expression in macrophages is regulated by PU.1 binding in human in publicly available ChIP-seq datasets. An increase in PU.1 binding in the mouse *Tspo* promoter after IFNγ treatment was observed (Fig. 1c). However, PU.1 binding to the human *TSPO* promoter was decreased after IFNγ treatment (Fig. 1d). To test whether the reduced PU.1 binding at the human *TSPO* promoter was due to reduced PU.1 expression, we analysed RNAseq data from the same set of samples. Expression of SPI-1, the gene that codes for PU.1, was not altered in human macrophages after IFNγ treatment (Fig. S1b), suggesting that the reduced binding of PU.1 to the human *TSPO* promoter region was unlikely to be due to reduced PU.1 levels. This suggests that repressive chromatin remodelling in the human cells leads to decreased PU.1 binding, a consequence of which could be the downregulation of *TSPO* transcript expression. This is consistent with the meta-analysis (Fig. 1a,b); although *TSPO* expression with inflammatory stimuli did not significantly change in most studies, in 8/9 (89%) of studies where TSPO did significantly change, it was downregulated (Fig. 1b). Together this data shows that *in vitro*, pro-inflammatory stimulation of mouse myeloid cells increases TSPO expression, histone marks in the enhancer regions and PU.1 binding. These changes are not found following pro-inflammatory stimulation of human myeloid cells.

### The presence of the AP1 binding site in the TSPO promoter and LPS inducible TSPO expression is unique to the *Muroidea* superfamily of rodents

To understand why TSPO expression is inducible by pro-inflammatory stimuli in mouse but not human myeloid cells, we performed multiple sequence alignment of the *TSPO* promoter region of 15 species including primates, rodents, and other mammals (Fig. 2). We found that an AP1 binding site is present uniquely in a subset of species within the *Muroidea* superfamily of rodents including mouse, rat and chinese hamster (Fig. 2a). These binding sites were not present in other rodents (squirrel, guinea pig), nor in other non-rodent mammals (Fig. 2a). We generated a phylogenetic tree which shows a clear branching in the *TSPO* promoter of rat, mouse and chinese hamster from the other rodents and non-rodent mammals (Fig. 2b). Differential motif enrichment analysis of the *TSPO* promotor region between *Muroidea* vs non-*Muroidea* species confirmed a significant enrichment of the AP1 binding site in the *Muroidea* promoter (Fig. 2c). We expanded this motif search and *TSPO* promoter sequence divergence analysis to a wider range of 24 rodent species from the *Muroidea* superfamily and other non*-Muroidea* rodents. Again, we found that the AP1 site is confined only to a subset of the superfamily *Muroidea* (Fig. S2).

**Figure 2.**
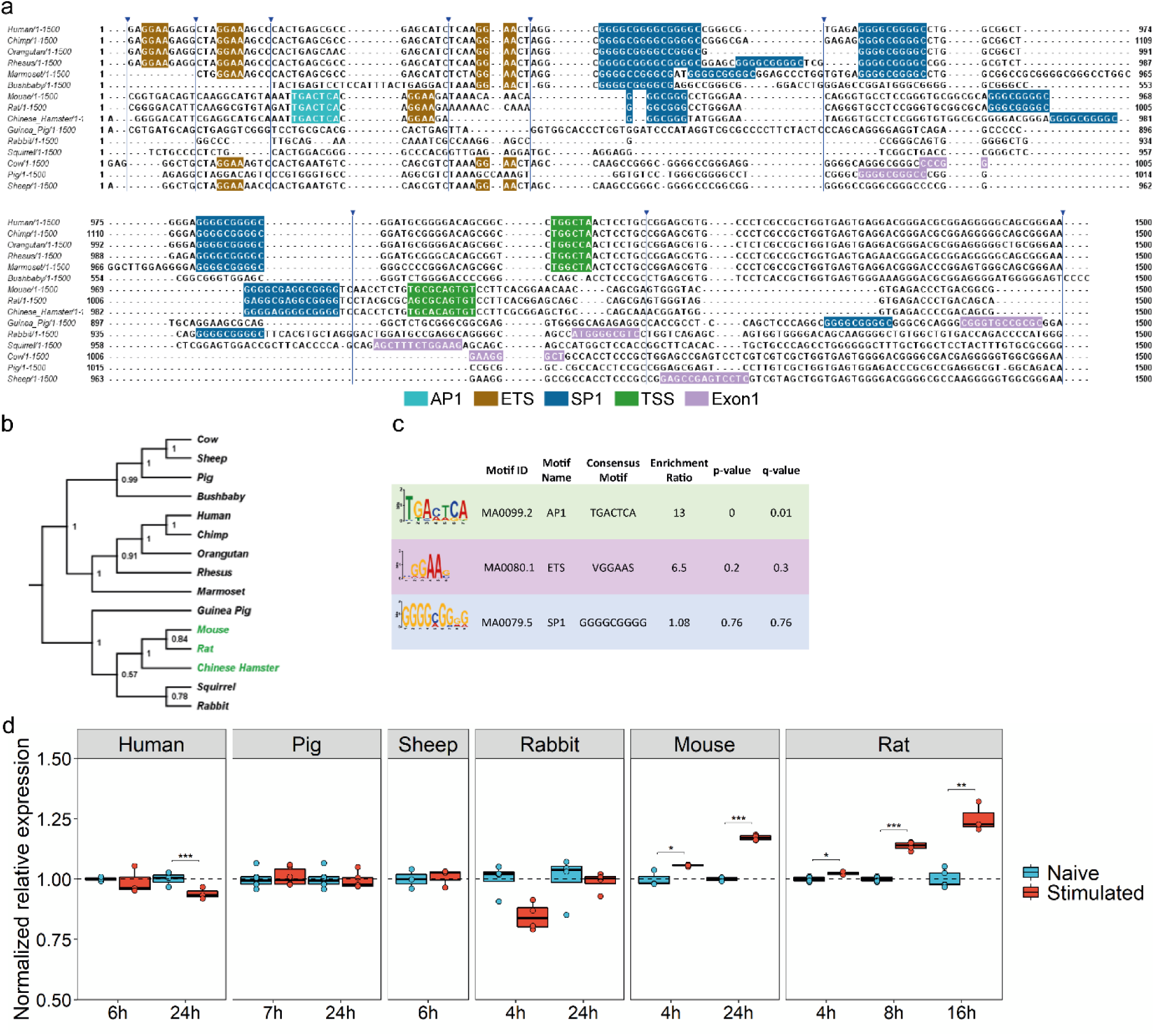
AP1 binding site in the TSPO promoter and LPS inducible TSPO expression is unique to the Muroidea superfamily of rodents. **a** Multiple sequence alignment of TSPO promoter region of 15 species from primate, rodent, non-primate mammals. AP1 (cyan) and an adjacent ETS (brown) site is present in only a sub-group of rodent family which includes mouse, rat and Chinese hamster. The ETS site which binds transcription factor PU.1 is present across species. SP1 (blue) site is found in the core promoter close to the TSS (green). For species where the TSS is not known Exon1 (pink) location is shown. Blue arrowhead indicates sequence without any motif hidden for visualization. **b** Phylogenetic tree is showing a clear branching of rat, mouse and Chinese hamster TSPO promoter from the rest of the species from rodents. Primates including marmoset forms a separate clade while sheep, cow and pig are part for the same branch. Green highlights represent species that contain the AP1 site in TSPO promoter. Phylogenetic tree was generated using the Maximum Parsimony method in MEGA11. The most parsimonious tree with length = 4279 is shown. The consistency index (CI) is 0.760458 (0.697014) and the retention index is 0.656386 (RI) (0.656386) for all sites and parsimony-informative sites (in parentheses). The percentage of replicate trees in which the associated taxa clustered together in the bootstrap test (1000 replicates) are shown next to the branches. d Differential motif enrichment analysis between rodent vs non-rodent TSPO promoter region by SEA tools from MEME-suite confirms the significant enrichment of AP1 site in rodent promoter whereas SP1 site does not show any differential enrichment. TSS; Transcription start site. **d** TSPO gene expression in macrophages or microglia isolated from multiple species after LPS stimulation. In line with the multiple sequence alignment of the TSPO promoter, species (mouse, rat) that contains an adjacent AP1 and ETS motif shows an upregulation of TSPO gene after LPS stimulation. Species lacking (human, pig, sheep, rabbit) those sites show a downregulation or no change in expression after stimulation.

Silencing AP1 impairs LPS induced TSPO expression in the immortalized mouse BV2 cell line^27^. We therefore tested the hypothesis that LPS inducible TSPO expression occurs only in species with the AP1 binding site in the promoter region. In species that lack the AP1 binding site (human, pig, sheep, rabbit), TSPO expression was not induced by LPS (Fig. 2d). However, in the rat, where the AP1 binding site is present, TSPO was increased under these conditions (Fig. 2d).

### Microglial TSPO expression is unchanged in the AD hippocampus, but is increased in amyloid mouse models

Microglia-neuronal interactions, which modulate microglia inflammatory phenotype^1^, are lost in monocultures *in vitro*. We therefore examined TSPO expression within inflammatory microglia *in situ* with quantitative neuropathology using *postmortem* samples from AD (Table S1). We compared data from human *postmortem* AD brain to the *App^NL-G-F^* and TAU^P301S^ mouse models.

We examined the hippocampal region, one of the most severely affected regions in AD^28, 29^, comparing it to non-neurological disease controls (Fig. 3a-c). No increases were observed in the number of IBA1+ microglia (Fig. 3d), HLA-DR+ microglia (Fig. 3e) or astrocytes (Fig. 3f) and the density of TSPO+ cells in AD did not differ compared to controls (Fig. 3g). Additionally, there was no increase in TSPO+ microglia (Fig. 3h,i) and astrocytes (Fig. 3j). We then quantified TSPO+ area (µm^2^) in microglia and astrocytes as an index of individual cellular expression (see methods). There was no difference in individual cellular TSPO expression in microglia (Fig. 3k) or astrocytes (Fig. 3k) in AD relative to controls.

**Figure 3.**
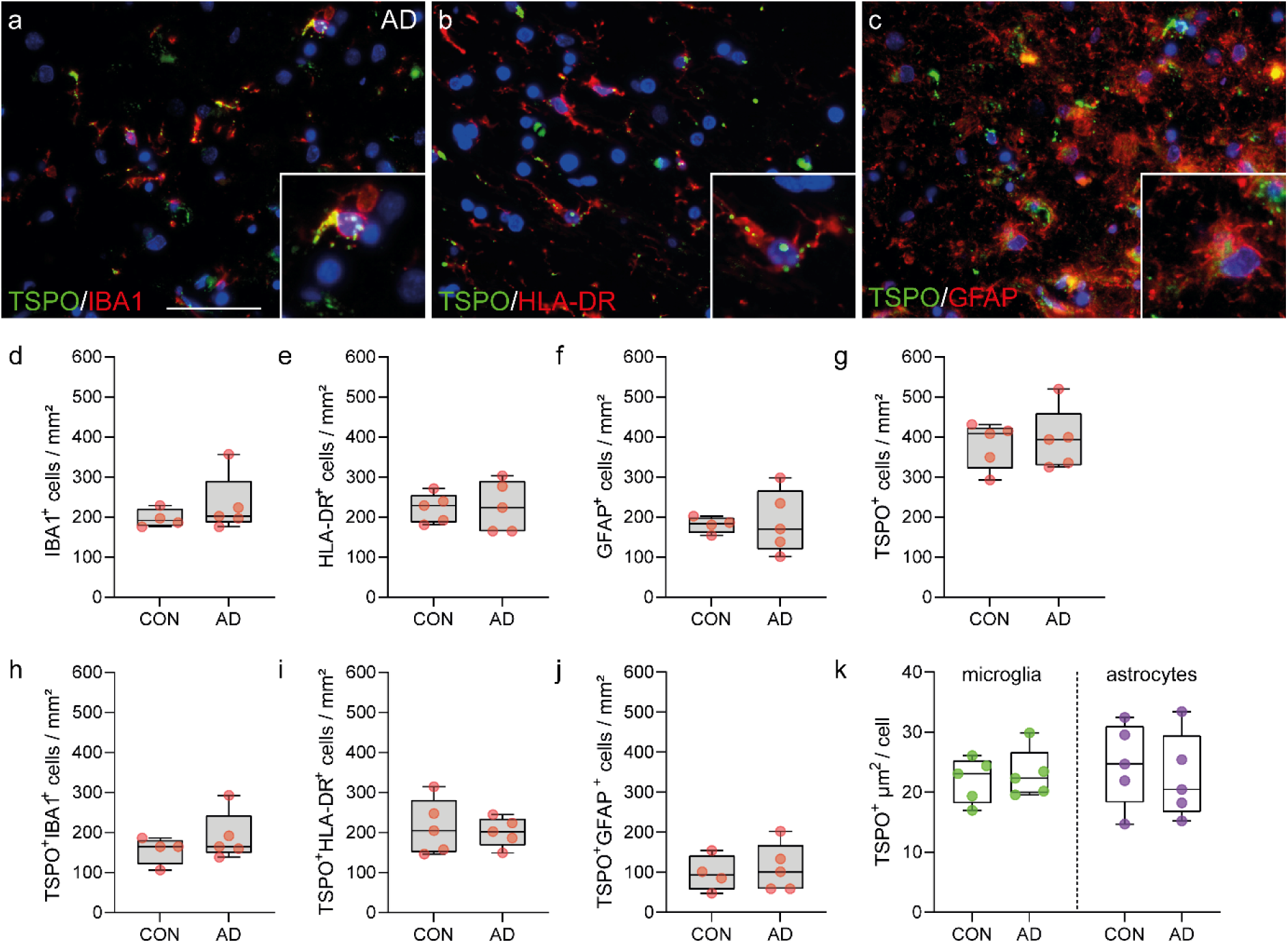
TSPO expression is not altered in the AD hippocampus. **a-c** Representative images of TSPO expression in microglia and astrocytes in AD hippocampus. **d-g** no increases were observed in microglia (P=0.5159, U=7, ranks=17, 28), activated microglia (P=0.8997, t=0.1301, df=8) astrocytes (P = 0.8599, t=0.1831, df=7) or TSPO+ cells (P = 0.7329, t=0.3534, df=8) in the AD hippocampus. **h-j** Concurrently no increases were observed in the number of TSPO+IBA1+ microglia (P = 0.3573, t=0.9854, df=7), TSPO+HLA-DR+ microglia (P = 0.7239, t=0.3659, df=8) and astrocytes (P = 0.7181, t=0.3760, df=7). **k** Even though microglia in the AD brain show signs of activation microglia do not upregulate TSPO expression in the hippocampus (P = 0.6717, t=0.4398, df=8), nor do astrocytes (P = 0.6475, t=0.4750, df=8). Statistical significance in **d-k** was determined by a two-tailed unpaired *t*-test or Mann-Whitney U-test when not normally distributed. Box and whiskers mark the 25^th^ to 75^th^ percentiles and min to max values, respectively, with the median indicated. Scale bar = 50µm, inserts are digitally zoomed in (200%).

We next conducted multiplexed proteomics with imaging mass cytometry (IMC) for further characterisation of cellular phenotype. As with the IHC, we did not see an increase in microglial density, as defined by the number of IBA1+ cells per mm^2^, (Fig. S3a) nor in the density of astrocytes (Fig. S3b). Furthermore, again in agreement with the IHC, we did not see an increase in the number of microglia and astrocytes expressing TSPO (Fig. S3c,d). However, IMC did reveal an increase in CD68+ microglia cells (Fig S3e) in AD compared to control, providing evidence, consistent with the literature^30, 31^, that microglia are activated in AD. However, despite microglial activation, we did not find an increase in individual cellular TSPO expression, defined here as mean cellular TSPO signal, in either microglia (Fig. S3f) or astrocytes (Fig. S3g) in AD donors relative to control. Because proximity to amyloid plaques is associated with activation of microglia^30^, we next tested whether cellular TSPO expression was higher in plaque microglia relative to (more distant) non-plaque microglia in the same tissue sections from the AD brains only. We saw no differences in cellular TSPO expression between the plaque and non-plaque microglia (Fig. S3h).

We next compared the human AD data to that from mouse *App^NL-G-F^* (Fig. 4a,b) and TAU^P301S^ (Fig. 4,i,j). The *App^NL-G-F^* model avoids artefacts introduced by APP overexpression by utilising a knock-in strategy to express human APP at wild-type levels and with appropriate cell-type and temporal specificity^18^. In this model, APP is not overexpressed. Instead, amyloid plaque density is elevated due to the combined effects of three mutations associated with familial AD (NL; Swedish, G: Arctic, F: Iberian). The *App^NL-G-F^* line is characterised by formation of amyloid plaques, microgliosis and astrocytosis^18^. We also investigated TSPO expression in a model of tauopathy, TAU^P301S^ mice, which develop tangle-like inclusions in the brain parenchyma associated with microgliosis and astrocytosis^19^. The use of these two models allows differentiation of effects of the amyloid plaques and neurofibrillary tangles on the expression of TSPO in the mouse hippocampus. In *App^NL-G-F^* mice, an increase in the density of microglia was observed at 28-weeks (Fig. 4c), but not in the density of astrocytes (Fig. 4d). An increase in TSPO+ cells was also observed (Fig. 4e), due to an increase in numbers of TSPO+ microglia and macrophages (Fig. 4f). No differences were observed in the density of TSPO+ astrocytes in *App^NL-G-F^* at 10 weeks, although a small (relative to that with microglia) increase was observed at 28 weeks (Fig. 4g). Finally, we then quantified TSPO+ area in microglia and astrocytes as an index of TSPO expression in individual cells. In contrast to the human data, expression of TSPO in individual cells was increased by 3-fold in microglia in the *App^NL-G-F^* mice at 28 weeks (Fig. 4h). It was unchanged in astrocytes. In the TAU^P301S^ mice, no differences were observed in microglia (Fig. 4k) or astrocyte (Fig. 4l) densities, in TSPO+ cell density (Fig. 4m), or in the density of TSPO+ microglia (Fig. 4n) or of TSPO+ astrocytes (Fig. 4o) in the hippocampus at either 8 or 20 weeks (Fig. 4) However, as with the *App^NL-G-F^* mouse (and in contrast to the human), a 2-fold increase in individual cellular TSPO expression was observed within microglia in TAU^P301S^ mice (Fig 4p). Again, as with the *App^NL-G-F^* mouse, individual cellular TSPO expression within astrocytes was unchanged.

**Figure 4.**
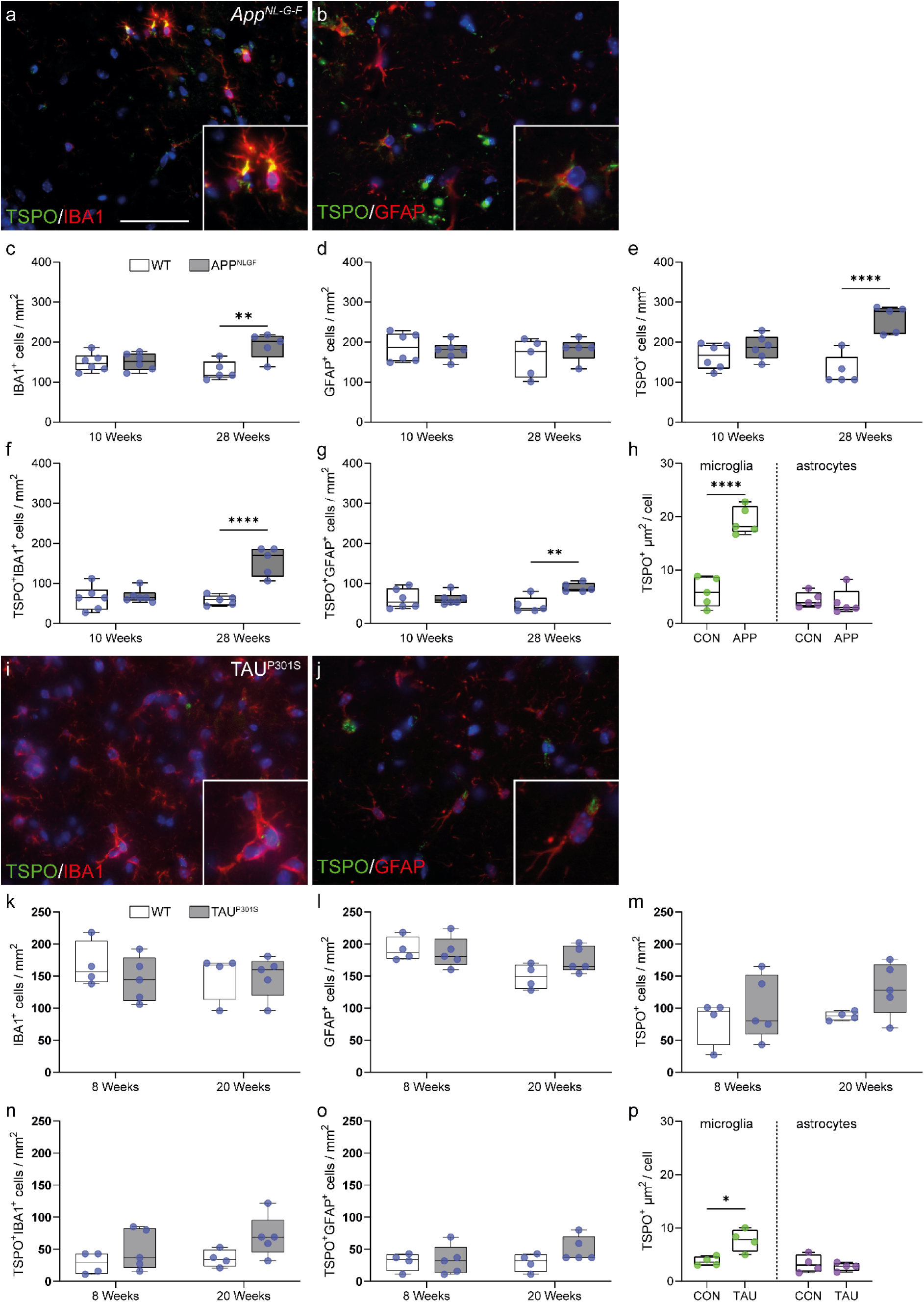
Microglia in the *App^NL-G-F^* and TAU^P301S^ model increase TSPO expression. **a,b** Representative images of TSPO expression in microglia and astrocytes in *App^NL-G-F^* hippocampus. **c** An increase was observed in IBA1+ microglia at 28 weeks (P = 0.0078, t=3.522, df=8) but not 10 weeks (P = 0.8788, t=0.1565, df=10) in *App^NL-G-F^* hippocampus compared to control. **d** No increase in astrocytes was observed (10 weeks: P = 0.6266, t=0.5019, df=10; 28 weeks: P = 0.4425, t=0.8080, df=8). **e** TSPO+ cells were increased at 28 weeks (P = 0.0079, U=0, ranks=15, 40) but not at 10 weeks (P = 0.2375, t=1.257, df=10) in the *App^NL-G-F^* mice. **f,g** Both TSPO+ microglia (P = 0.0005, t=5.658, df=8) and astrocytes (P = 0.0030, t=4.207, df=8) were increased at 28 weeks in the hippocampus of *App^NL-G-F^* mice but not at 10 weeks (microglia: P = 0.7213, t=0.3670, df=10; astrocytes: P = 0.9561, t=0.056, df=10) . **h** Activated microglia (P < 0.0001, t=7.925, df=8), but not astrocytes (P = 0.3095, U=7, ranks=33, 22), in the *App^NL-G-F^* model have increased TSPO expression. **i,j** Representative images of TSPO expression in microglia and astrocytes in TAU^P301S^ hippocampus. **k-m** No increases in microglia (8 weeks: P = 0.3687, t=0.9608, df=7; 20 weeks; P = 0.9647, t=0.04580, df=7), astrocytes (8 weeks: P = 0.7353, t=0.3519, df=7; 20 weeks; P = 0.0870, t=1.989, df=7) or TSPO+ cells (8 weeks: P = 0.8492, U=9, ranks=19, 26; 20 weeks; P = 0.0876, t=1.985, df=7) were observed in the hippocampus of TAU^P301S^ mice. **n,o** No increase was observed in the number of TSPO+ microglia (8 weeks: P = 0.2787, t=1.174, df=7; 20 weeks; P = 0.0907, t=1.961, df=7) or astrocytes (8 weeks: P = 0.8684, t=0.1718, df=7; 20 weeks; P = 0.1984, U=4.5, ranks=14.5, 30.5). **p** Microglia in the TAU^P301S^ increase TSPO expression (P = 0.0133, t=3.471, df=6) whereas astrocytes do not (P = 0.5800, t=0.5849, df=6). Statistical significance in **c-h** and **k-p** was determined by a two-tailed unpaired *t*-test or Mann-Whitney U-test when not normally distributed. Box and whiskers mark the 25th to 75th percentiles and min to max values, respectively, with the median indicated. Scale bar = 50µm, inserts are digitally zoomed in (200%).

In summary, we showed that TSPO cellular expression is increased within microglia from *App^NL-G-F^* and TAU^P301S^ mice, but not in microglia from AD tissue. TSPO was also unchanged in astrocytes from both mouse models and the human disease.

### Microglial TSPO is upregulated in SOD1^G93A^ mice but not in ALS

Spinal cord and brain microglia differ with respect to development, phenotype and function^32^. We therefore next investigated ALS (Table S2), that primarily affects the spinal cord rather than the brain. We compared this data to that from the commonly used SOD1^G93A^ mouse model of ALS. TSPO expression was investigated in the ventral horn and lateral columns of the spinal cord in cervical, thoracic, and lumbar regions (Fig. 5a-c). An increase in microglia (Fig. 5d), HLA-DR+ microglia (Fig. 5e) and astrocytes (Fig. 5f) was observed in human ALS spinal cord. The density of TSPO+ cells was increased by 2.5-fold in ALS spinal cords across all regions when compared to controls (Fig. 5g). No additional changes were found when stratifying the cohort based on disease duration or spinal cord regions, white or grey matter, or spinal cord levels. In comparison to the controls, ALS samples exhibited a 3-fold increase in the density of TSPO+ microglia (TSPO+IBA1+ cells, Fig. 5h) and a 3-fold increase in TSPO+ activated microglia/macrophages (TSPO+HLA-DR+ cells, Fig. 5i). A 2.5-fold increase in the density of TSPO+ astrocytes (TSPO+GFAP+ cells) was observed in ALS compared to control (Fig. 5j). We then quantified TSPO+ area in microglia and astrocytes as an index of individual cellular TSPO expression (Fig. 5k). No increase in TSPO+ area (µm^2^) was found in microglia or astrocytes in ALS when compared to control (Fig. 5k), implying that TSPO expression does not increase in microglia or astrocytes with ALS.

**Figure 5.**
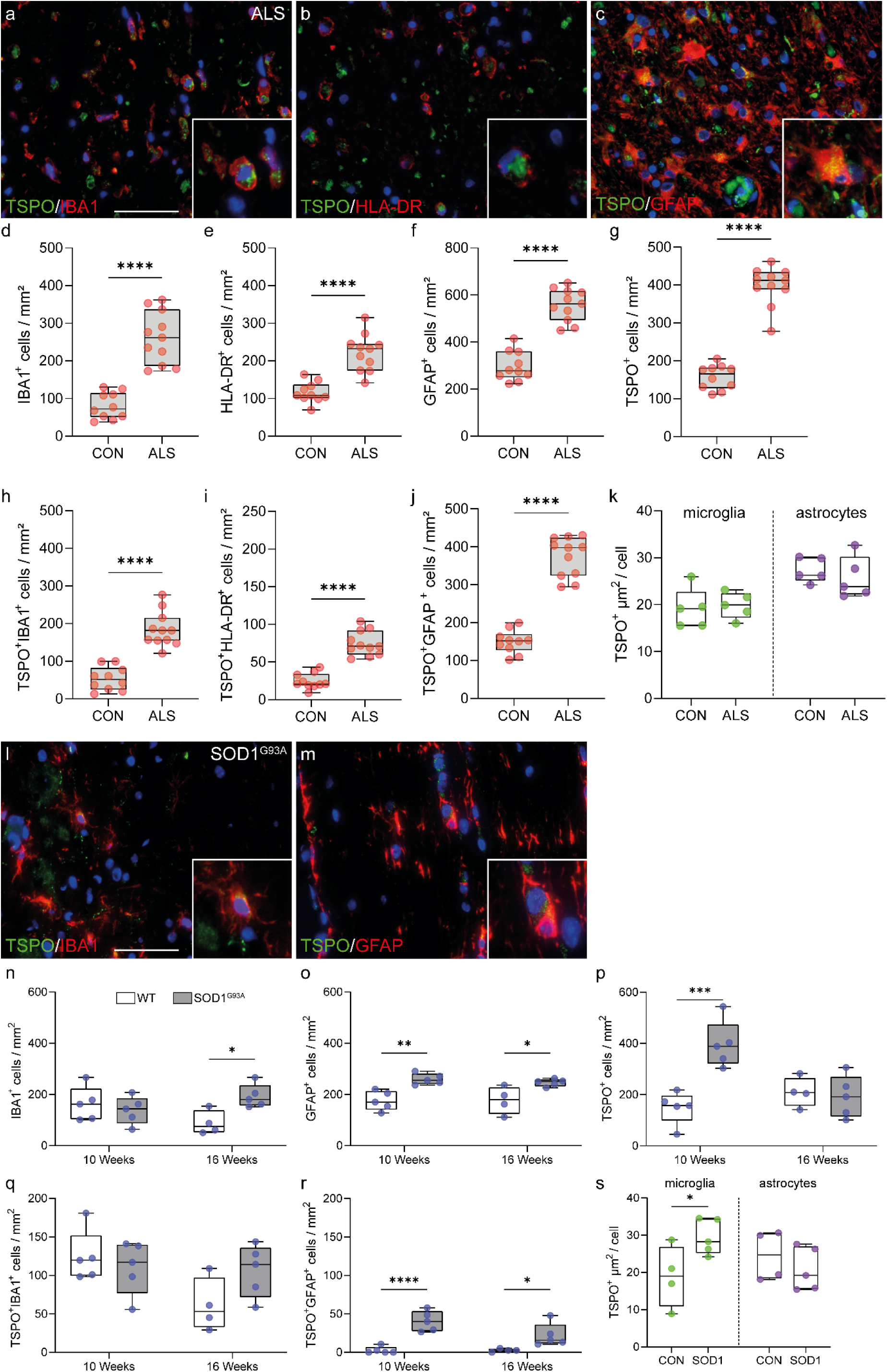
TSPO is increased in microglia in SOD1^G93A^ mice but not in ALS spinal cord. **a-c** Representative images of TSPO expression in microglia and astrocytes in ALS spinal cord. **d-f** An increase was observed in microglia (P < 0.0001, t=7.445, df=19), HLA-DR+ microglia (P < 0.0001, t=6.007, df=19), and astrocytes (P < 0.0001, t=9.024, df=19) in ALS spinal cord when compared to controls. **g** A 2.5-fold increase of TSPO+ cells (P < 0.0001, t=12.88, df=19) was observed in the ALS spinal cord. **h,i** Up to a 3.4-fold increase in the density of TSPO+ microglia (TSPO+IBA1+ cell, P < 0.0001, t=7.541, df=19) (TSPO+HLA-DR+ cells, P < 0.0001, t=3.368, df=19) was observed. **j** TSPO+ astrocytes were significantly increased (P < 0.0001, t=11.77, df=19) in the spinal cord of ALS patients. **k** The increase in activated microglia and astrocytes was not associated with an increase in TSPO expression in microglia (P = 0.7684, t=0.3046, df=8) or in astrocytes (P = 0.5047, t=0.6985, df=8). **l,m** Representative images of TSPO expression in microglia and astrocytes in SOD1^G93A^ spinal cord. **n** An increase was observed in microglia in SOD1^G93A^ spinal cord when compared to controls at 16 weeks (P=0.0115, t=3.395, df=7) but not at 10 weeks (P = 0.5334, t=0.6509, df=8). **o** An increase for astrocytes was observed for both 10 weeks (P = 0.0024, t=4.362, df=8) and 16 weeks (P = 0.0248, t=2.848, df=7) **p** An increase in TSPO+ cells was observed at 10 weeks (P = 0.0011, t=4.931, df=8) but not 16 weeks (P = 0.7299, t=0.3594, df=7). **q** No increase in the number of TSPO+ microglia was observed (10 weeks: P = 0.5244, t=0.6656, df=8; 16 weeks, P = 0.0930, t=1.944, df=7). **r** TSPO+ astrocytes were increased up to 15-fold in the spinal cord of SOD1^G93A^ mice (10 weeks: P = 0.0003, t=6.085, df=8; 16 weeks: P = 0.382, t=2.548, df=7). **s** Despite no increase in the number of TSPO+ microglia, an increase in the amount of TSPO per cell was observed in microglia (P = 0.0451, t=2.435, df=7), but not astrocytes (P = 0.4052, t=0.8856, df=7). Statistical significance in **d-k**, and **o-s** was determined by a two-tailed unpaired *t*-test. Box and whiskers mark the 25^th^ to 75^th^ percentiles and min to max values, respectively, with the median indicated. Scale bar = 50µm, inserts are digitally zoomed in (200%).

SOD1^G93A^ mice express high levels of mutant SOD1 that initiates adult-onset neurodegeneration of spinal cord motor neurons leading to paralysis, and as such these mice have been used as a preclinical model for ALS^20^. To determine the extent to which TSPO+ cells were present in SOD1^G93A^ mice TSPO+ microglia and astrocytes were quantified with immunohistochemistry in the white and grey matter of the spinal cord (Fig. 5l,m). An increase was observed in the total number of microglia (Fig. 5n) and astrocytes (Fig. 5o) in 16-week old SOD1^G93A^ mice but not in 10 week old animals (Fig. 6c,d). The density of TSPO+ cells was increased 2- to 3-fold in presymptomatic disease (10 weeks) compared to non-transgenic littermate control mice in both white and grey matter (Fig. 5p). Increases in the density of TSPO+IBA+ cells were not observed in SOD1^G93A^ mice compared to control animals (Fig. 5q). However, a significant 8- to 15-fold increase in the density of TSPO+GFAP+ astrocytes was observed in 10- and 16-week old SOD1^G93A^ mice compared to 10- and 16-week old wild-type mice (Fig. 5r). Finally, we then quantified TSPO+ area in microglia and astrocytes as an index of individual cellular TSPO expression. In contrast to the human data, where there was no change in disease samples relative to controls, expression of TSPO in individual cells was increased by 1.5-fold in microglia in the rodent model. As with the *App^NL-G-F^* and TAU^P301S^ mice above, TSPO expression within astrocytes was unchanged (Fig. 5s).

**Figure 6.**
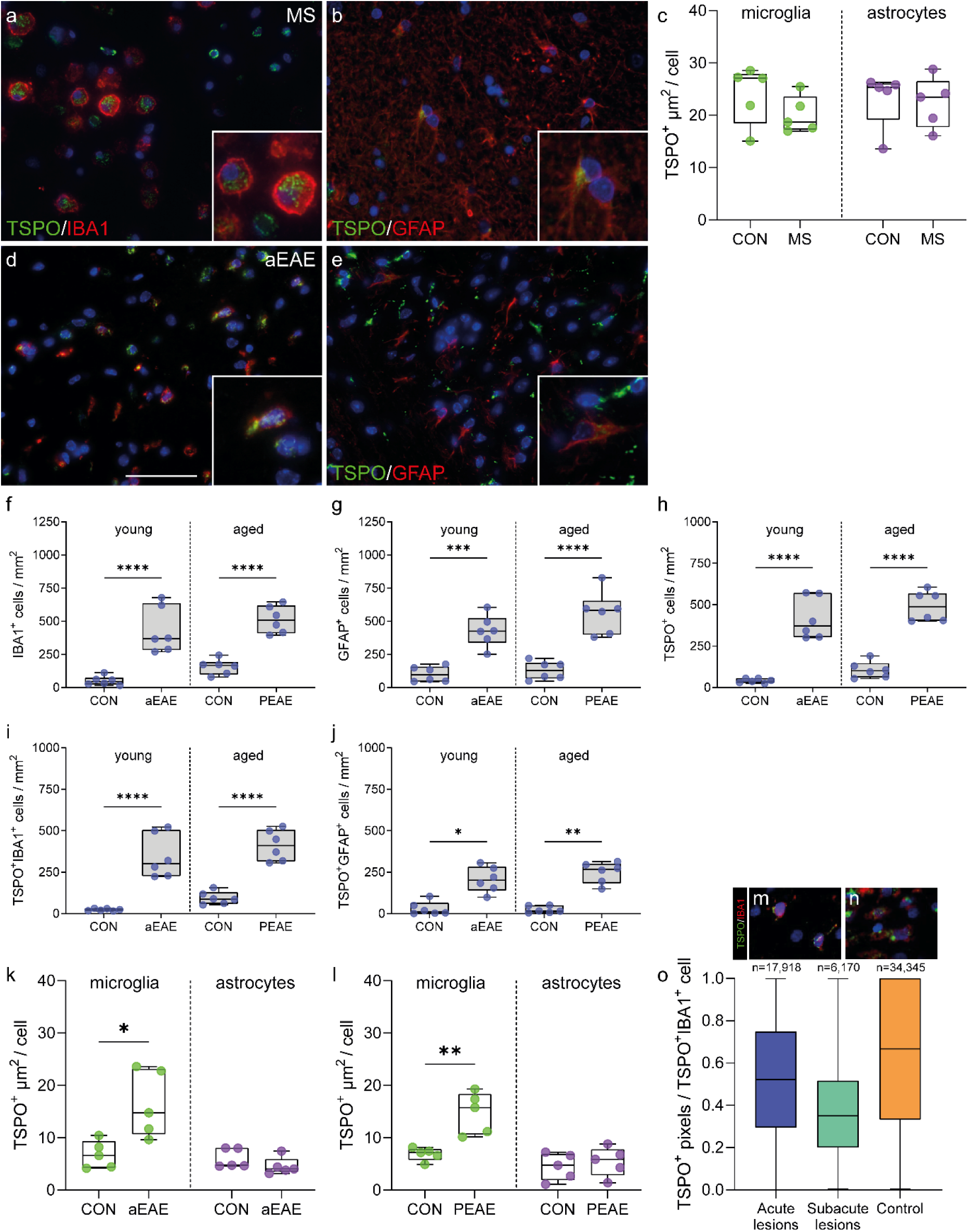
Microglia in mouse aEAE and PEAE, and marmoset EAE, but not MS, increase TSPO expression. **a,b** Representative images of TSPO+ microglia and astrocytes in MS. **c** TSPO+ microglia (P = 0.2278, t=1.306, df=8) and astrocytes (P = 0.5476, U=9, ranks=31, 24) do not increase TSPO expression in MS. **d,e** Representative images of TSPO expression in microglia and astrocytes in EAE mice. **f-h** microglia (P < 0.0001, F_(3,20)_=25.68), astrocyte (P < 0.0001, F_(3,20)_=25.51), and TSPO+ cell numbers (P < 0.0001, F_(3,20)_=44.53), are increased during disease in aEAE mice and PEAE. **i,j** An increase in both TSPO+ microglia (P < 0.0001, F_(3,20)_=30.93) and TSPO+ astrocytes (P = 0.0005, K-W=17.72) is observed during disease. **k,l** TSPO+ microglia increase TSPO expression in aEAE mice (P = 0.0136, t=3.152, df=8), and in PEAE mice (P = 0.0028, t=4.248, df=8). Astrocytes do not increase TSPO expression in aEAE (P = 0.0556, U=3, ranks=37, 18), and PEAE (P = 0.5918, t=0.5584, df=8). **m,n** Representative images of TSPO+ microglia in marmoset EAE. **o** TSPO+ pixels are not increased in acute and subacute lesions in marmoset EAE relative to control. Statistical significance in **f-j,o** was determined by a one way ANOVA or Kruskal-Wallis test when not normally distributed, and by a two-tailed unpaired *t*-test or Mann-Whitney U-test when not normally distributed in **c,k** and **l**. Holm-Sidak’s and Dunn’s multiple comparisons were performed. Box and whiskers mark the 25^th^ to 75^th^ percentiles and min to max values, respectively, with the median indicated. Scale bar = 50µm, inserts are digitally zoomed in (200%).

In summary, consistent with the data from AD and relevant mouse models, we have shown that TSPO expression is increased within microglia from SOD1^G93A^ mice, but not increased in microglia from human ALS tissue. TSPO also was unchanged in astrocytes from the SOD1^G93A^ mice and the human disease relatively to those in the healthy control tissues.

### Increased myeloid cell TSPO expression is found in mouse EAE, but not in MS or marmoset EAE

Having found no evidence of increased TSPO expression in activated microglia in human neurodegenerative diseases affecting the brain or spinal cord, we next examined MS as an example of a classical neuroinflammatory disease characterised by microglia with a highly activated pro-inflammatory phenotype. We compared data from human *postmortem* MS brain (Table S3) to mice with EAE (Table S4). We also examined brain tissue from marmoset EAE (Table S5), as *antemortem* MRI assessments in these animals allow for identification of acute lesions which are highly inflammatory.

We previously defined TSPO cellular expression in MS^16, 33^. HLA-DR+ microglia expressing TSPO were increased up to 14-fold in active lesions compared to control^33^, and these microglia colocalised with CD68 and had lost homeostatic markers P2RY12 and TMEM119, indicating an activated microglial state^16^. Here we quantified individual cellular TSPO expression in both microglia and astrocytes by comparing cells in active white matter lesions to white matter from control subjects. Consistent with the human data from AD and ALS, there was no difference in TSPO expression in individual microglia or astrocytes in MS compared to control tissue (Fig. 6a-c).

We next investigated the relative levels of TSPO expression (Fig. 6d-l) in microglia and astrocytes in acute EAE (aEAE), a commonly used experimental mouse model of MS^21, 34^. Neurodegenerative diseases typically occur in old age, whereas aEAE and the AD and ALS relevant rodent models described above are induced in young mice. As age might affect TSPO regulation^35^, we also investigated TSPO expression in progressive EAE (PEAE), a model where the pathology is induced in aged mice (12 months).

Increases in numbers of both microglia and astrocytes were observed in aEAE as well as in PEAE mice compared to their respective young and old control groups (Fig. 6f,g). Similarly, increases were observed in the number of TSPO+ microglia and TSPO+ astrocytes in both aEAE and PEAE relative to their respective controls (Fig. 6h-j). When comparing the young control mice (aEAE controls) with the old control mice (PEAE controls), no differences were observed in microglial and TSPO+ microglial density (Fig. 6f,i). Similarly, there was no difference in density of astrocytes or TSPO+ astrocytes between these two control groups (Fig. 6g,j).

To investigate individual cellular TSPO expression, TSPO+ area was measured in microglia and astrocytes. Individual microglia expressed 3-fold greater TSPO and 2-fold greater TSPO in aEAE and PEAE respectively, relative to their control groups. The individual cellular TSPO expression was not higher in microglia from young mice relative to old mice. Again, as with the SOD1^G93A^, *App^NL-G-F^*, and TAU^P301S^ mice, individual cellular TSPO expression within astrocytes was unchanged.

Finally, we investigated TSPO expression in EAE induced in the common marmoset (Callithus jacchus)(Fig. S4, Fig. 6m-o), a non-human primate which, like humans, lacks the AP1 binding site in the core promoter region of TSPO. Both the neural architecture and the immune system of the marmoset are more similar to humans than are those of the mouse^36–38^. Marmoset EAE therefore has features of the human disease which are not seen in mouse EAE, such as perivenular white matter lesions identifiable by MRI, B cell infiltration and CD8+ T cell involvement. Marmosets were scanned with MRI biweekly, which allowed the ages of lesions to be determined and the identification of acute lesions including pro-inflammatory microglia. In acute and subacute lesions, there was an increase of up to 27-fold in the density of TSPO+ microglia relative to control (Fig. S4a-c) and these microglia bore the hallmarks of pro-inflammatory activation. However, TSPO expression in individual microglia, here defined as the percentage of TSPO^+^ pixels using immunofluorescence, was not increased in acute or subacute lesions relative to control (Fig. 6o).

In summary, and consistent with the AD and ALS data, we have shown that individual cellular TSPO expression is increased in microglia in EAE in both young and aged mouse models, but it is not increased in microglia from MS lesions nor marmoset EAE acute lesions. Again, consistent with previous data, astrocytes did not show an increase in TSPO expression in either MS or EAE.

### Single cell RNAseq shows *TSPO* gene expression is upregulated in activated mouse microglia, but not in activated human microglia

Methods for protein quantification by immunohistochemistry in *postmortem* brain are semiquantitative and therefore we also assessed *ex vivo* species-specific TSPO gene expression of microglial under pro-inflammatory conditions to add further confidence to our findings. We employed publicly available human and mouse scRNAseq datasets^39–44^. We first examined evidence for a pro-inflammatory microglial phenotype by quantifying the differential expression of homeostatic and/or activation markers. We then quantified the differential expression of TSPO in pro-inflammatory activated microglia using MAST^45^.

In a model of LPS exposure in the mouse^39^, scRNAseq yielded 2019 microglial cells that showed evidence of pro-inflammatory activation including a downregulation of the homeostatic marker *P2ry12* and an upregulation of activation markers *Fth1* and *Cd74* (Fig. 7a). In this population, *TSPO* was significantly upregulated. In a mouse model of acute EAE^40^, scRNAseq yielded 8470 pro-inflammatory activated microglial cells that showed significant downregulation of *P2ry12*, and a significant upregulation of *Fth1* and *Cd74* (Fig. 7b). *TSPO* was significantly upregulated. Finally, in the 5XFAD mouse model of AD^41^, scRNAseq yielded over 6203 microglial cells. Among them, 223 showed enrichment in disease-associated microglia (DAM) markers^41^, including increased expression of *Apoe*, *Trem2*, *Tyrobp* and *Cst7* (Fig. 7c). Compared to non-DAM cells, DAM cells showed a significant upregulation of *TSPO*.

**Figure 7.**
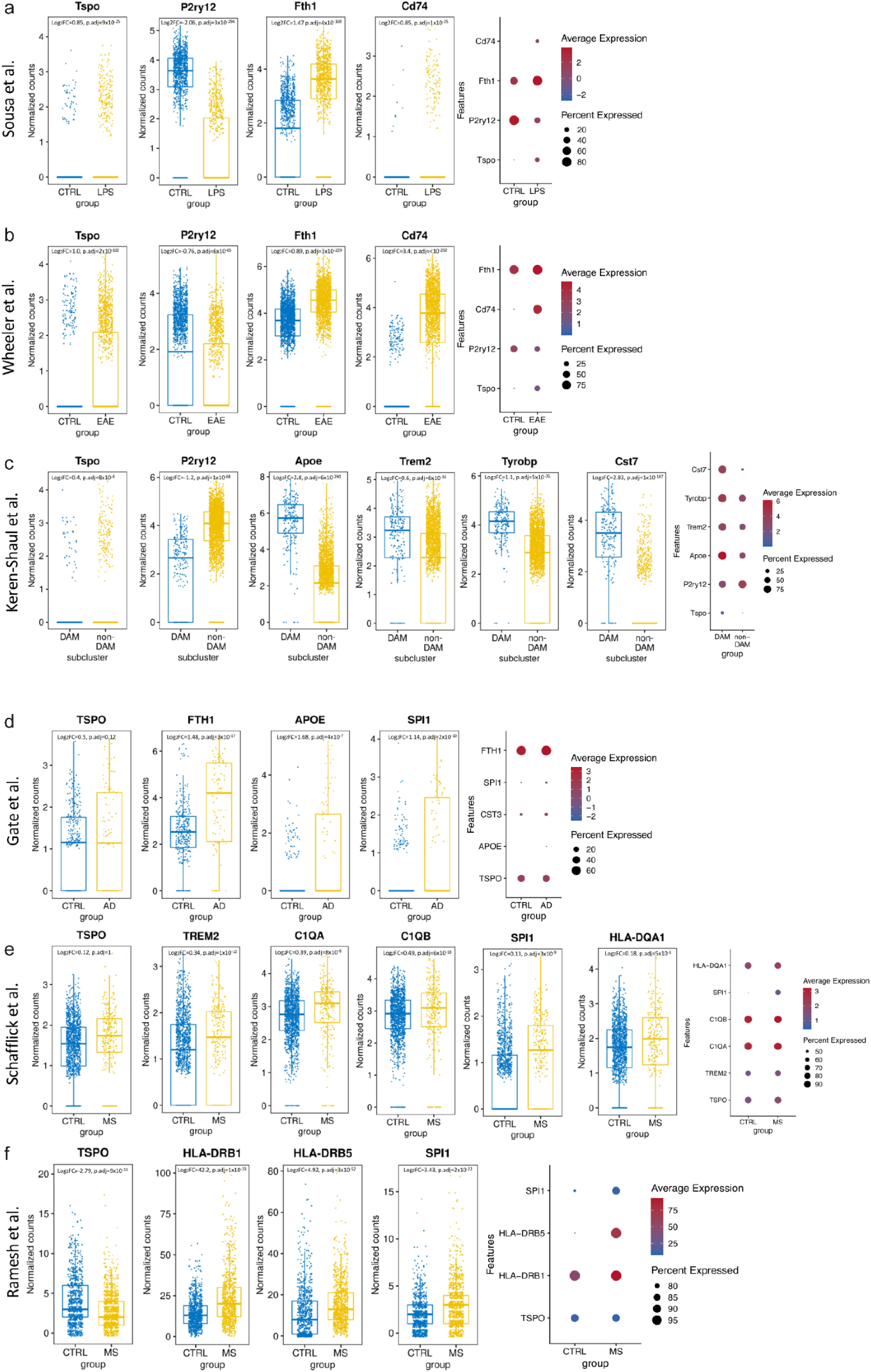
TSPO is increased in mouse but not human pro-inflammatory activated and disease-associated microglia. **a-c** Boxplots and dotplots showing the significantly elevated expression of *Tspo* in mouse models of pro-inflammatory activation using LPS (GSE115571), of acute EAE (GSE130119) and of AD (GSE98969). The percentage of cells that express *Tspo* in mouse microglia is relatively low, but it is considerably increased after LPS treatment, in the EAE model and in the DAM cells. **d-f** TSPO is not significantly upregulated in microglia-like cells from the CSF of AD (GSE134578) and MS (GSE138266) patients. The percentage of cells that express a given gene corresponds to the size of the dot, whereas the average expression corresponds to the fill colour of the dot.

In cerebrospinal fluid (CSF)-derived cells isolated from people with AD^42^, microglia-like cells (n=522) had an activated phenotype with a significant upregulation of *APOE*, *FTH1* and *SPI1* relative to controls. However, *TSPO* was not differentially expressed (Fig. 7d). In CSF isolated from people with MS^43^, microglia-like cells (n=1650) showed evidence of activation: *TREM2, C1QA, C1QB, SPI1, and HLA-DQA1* all were significantly upregulated^43^. However, TSPO was not differentially expressed in these cells (Fig. 7e). In a similarly designed study also using CSF-derived cells, microglia showing upregulation of *HLA-DRB1, HLA-DRB5* and *SPI1* also downregulated TSPO^44^ (Fig. 7f).

These experiments are consistent at the gene expression level with our own data at the protein expression level showing that the *TSPO* gene is not increased in microglia in AD or EAE, but is increased in their respective commonly used mouse models.

### TSPO is mechanistically linked to classical pro-inflammatory myeloid cell function in mice but not humans

Having demonstrated species-specific differences in TSPO expression and regulation, we then sought to examine TSPO function in mouse and human myeloid cells. We first examined the effect of pharmacological modulation of the classical microglial pro-inflammatory phenotype using the high affinity TSPO ligand, XBD173. Consistent with the literature^11–13^, we found that in primary mouse macrophages and the BV2 mouse microglial cell line, XBD173 reduced LPS induced release of proinflammatory cytokines (Fig. 8a,b,c). However, in primary human macrophages and in human induced pluripotent stem cell (hIPSC) derived microglia, XBD173 had no impact on the release of these cytokines, even at high concentrations associated with 98% TSPO binding site occupancy (Fig. 8d,e,f,g). We found similar results for zymosan phagocytosis. Primary mouse microglia demonstrated a dose dependant increase in phagocytosis upon exposure to XBD173 (Fig. 8h). However, we saw no increase in phagocytosis in primary human macrophages upon XBD173 exposure (Fig. 8i).

**Figure 8.**
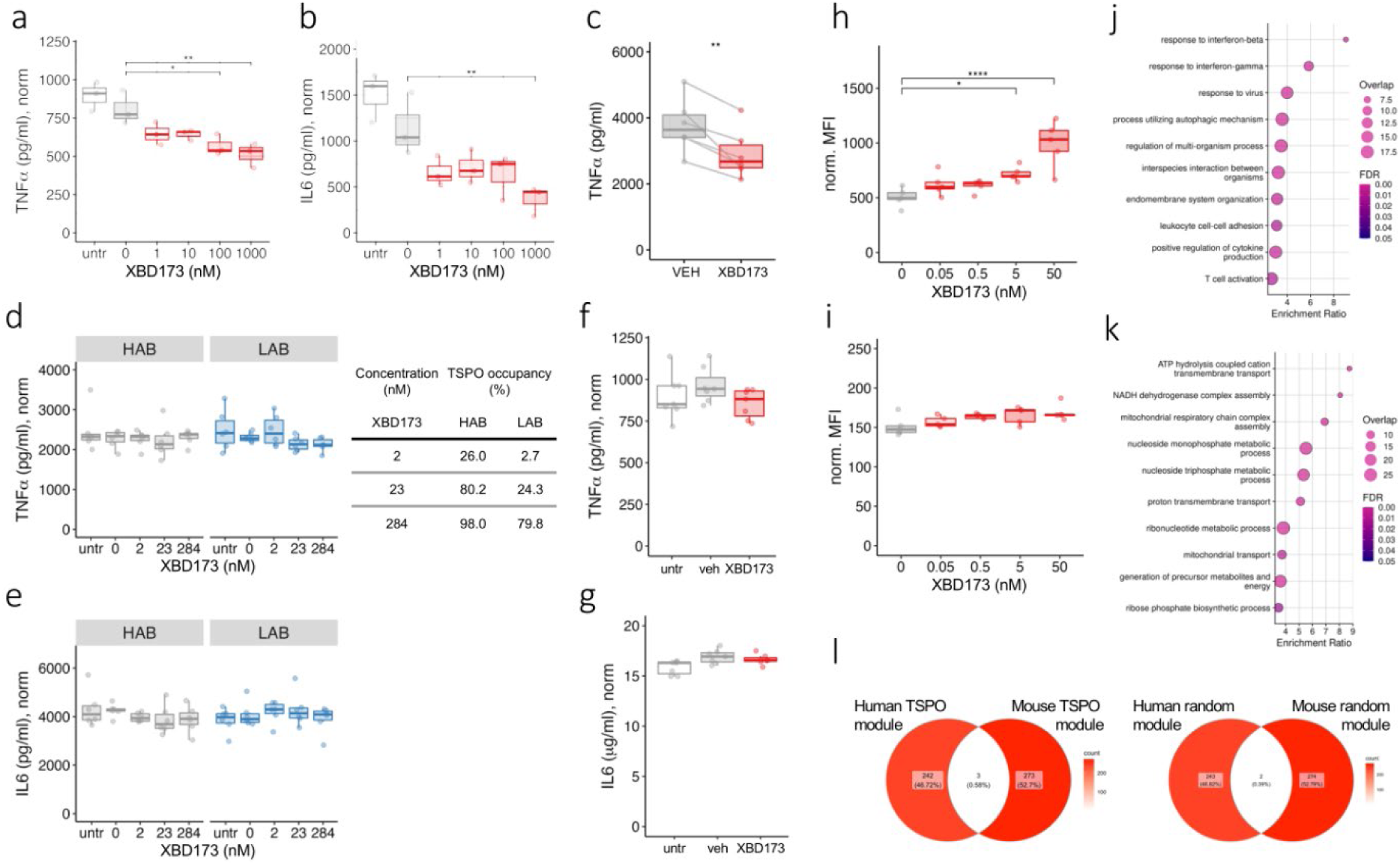
TSPO ligand XBD-173 modulates classical pro-inflammatory myeloid cell function in mouse but not human myeloid cells. **a-c.** The specific TSPO ligand XBD-173 reduces LPS-induced cytokine secretion in mouse BV2 microglia (**a,b**) and primary bone-marrow derived macrophages (**c**; BMDM, XBD = 10nM). (**a** P = 0.0007, F = 9.646, df = 5, n = 3, padj_(100)_ = 0.014, padj_(1000)_ = 0.003; **b** P = 0.0008, F = 9.282, df = 5, n = 3, padj_(1000)_ = 0.006; **c** P = 0.005, n = 6). **d-g** XBD-173 does not reduce LPS-induced cytokine secretion by human primary monocyte-derived macrophages derived from rs6971 AA individuals (high affinity binders, HAB), rs6971 TT individuals (low affinity binders, LAB) **(d,e)** or by hiPSC derived microglia-like cells **(f,g)** (**d** HAB: P = 0.8333, K-W = 1.4624, df = 4, n = 6; LAB: P = 0.141, K-W = 5.8624, df = 4, n = 6; **e** HAB: P = 0.09999, K-W = 7.7796, df = 4, n = 6, LAB: P = 0.2097, F = 0.68, df = 4, n = 6; **f** P = 0.057, n = 7, XBD = 200nM; **g** P = 0.423, n = 7, XBD = 200nM). **h,i** XBD-173 enhances phagocytosis in mouse BMDM (**h**) but not human monocytes (**i**) (**h** P < 0.0001, F = 12.07, df = 4, n = 5; **i** P = 0.1728, K-W = 6.376, df = 4, n = 5). **j-k** TSPO gene co-expression modules from naïve and pro-inflammatory primary macrophages in mouse and human. Gene ontology biological processes for the mouse TSPO module is enriched in classical proinflammatory pathways (**j**) and the human TSPO module is enriched for bioenergetic pathways (**k**). 3 genes overlap between mouse and human TSPO modules (**l**, left panel), compared to 2 genes overlapping between human and mouse random modules of the same size (**l**, right panel). Statistical significance in **a,b, d,e** and **i,j** was determined by one way ANOVA or Kruskal-Wallis test when not normally distributed and by a two-tailed unpaired *t*-test or Mann-Whitney U-test when not normally distributed in **c,f** and **g**. Box and whiskers mark the 25th to 75th percentiles and min to max values, respectively, with the median indicated.

XBD173 is metabolised by CYP3A4, which is expressed in myeloid cells. We therefore used LC-MSMS to quantify XBD173 in the supernatant in order to test the hypothesis that the lack of drug effect on human myeloid cells was due to depletion of XBD173. The measured concentration of XBD173 in the supernatant at the end of the assay was no different to the planned concentration (Fig. S5), excluding the possibility that XBD173 metabolism explained the lack of effect.

To understand if TSPO is associated with divergent functional modules in mouse and human we then used weighted gene co-expression network analysis to examine the genes whose expression are correlated with *TSPO* in mouse and human myeloid cells. To construct the gene co-expression networks, we used four publicly available and one in-house RNA-seq data from human (n = 47) and five publicly available mouse (n = 35) datasets of myeloid cells treated with LPS or LPS and IFNγ. In mouse myeloid cells, the gene ontology biological processes associated with the TSPO network related to classical pro-inflammatory functions such as responses to type 1 and 2 interferons, viruses and regulation of cytokine production (Fig 8j, Supplementary File 1). However, in human myeloid cells, the processes associated with the TSPO co-expression network related to bioenergetic functions such as ATP hydrolysis, respiratory chain complex assembly, and proton transport (Fig 8k, Supplementary File 1). There was no overlap in the genes that TSPO is co-expressed with in mouse, relative to human, myeloid cells (Fig 8l).

## Discussion

Microglial activation accompanies and is a major contributor to neurodegenerative and neuroinflammatory diseases^1, 4-6, 46^. A better understanding of microglial activation in combination with a technique that could reliably quantify activated microglia in the human brain would have broad utility to monitor disease progression as well as response to therapy. TSPO PET has been applied by many with this objective^9, 10^. Here we have tested the widely held assumption that *TSPO* cellular expression increases upon microglial activation. We examined *in vitro* data from isolated myeloid cells across 6 species, multiple sequence alignment of the TSPO promoter region across 34 species, and *ex vivo* neuropathological and scRNAseq data from human neuroinflammatory and neurodegenerative diseases, with relevant marmoset and young and aged mouse models. We show that TSPO expression increases in mouse and rat microglia when they are activated by a range of stimuli, but that this phenomenon is unique to microglia from a subset of species from the *Muroidea* superfamily of rodents. The increase in TSPO expression is likely dependant on the AP1 binding site in the core promoter region of TSPO. Finally, we showed that TSPO is mechanistically linked to classical pro-inflammatory myeloid cell function in mice but not humans.

This finding fundamentally alters the way in which the TSPO PET signal is interpreted, because it implies that the microglial component of the TSPO PET signal reflects density only, rather than a composite of density and activation phenotype. For example, in Parkinson’s Disease (PD) there is evidence of activated microglia in the *postmortem* brain but minimal change in microglial density^47^. Three well designed studies using modern TSPO radiotracers found no difference in TSPO signal between PD and controls groups^48–, 50^. The lack of increase in the TSPO PET signal is consistent with the data presented here, and should therefore not be interpreted as evidence for lack of microglial activation in PD.

Our study has several limitations. First, we have only examined microglia under certain pro-inflammatory conditions and cannot exclude the possibility that other stimulation paradigms would increase TSPO in human myeloid cells. However, the *in vitro* stimuli which were examined included a broad range of pro-inflammatory triggers, and the three human diseases are diverse with respect to the mechanisms underlying the activation of microglia. Second, the measurements of cellular TSPO expression we used in brain tissue are semi-quantitative. However, the same IHC quantification methods were used in all human and mouse comparisons, and these methods consistently detected cellular TSPO increases in mouse microglia despite not detecting analogous changes in human microglia. Furthermore, where IMC and immunofluorescence were used, the quantitative data were consistent with IHC. The neuropathology protein quantification was also consistent with gene expression measured by scRNAseq. Third, for RNAseq analysis, we were restricted to single cell rather than single nucleus experiments. This is because *TSPO* is detected in only 5-12% of microglial nuclei^51–54^ but ∼80% of microglial cells^39–44^. Fourth, the *in vitro* assay which most closely mimics *in vivo* PET data is radioligand binding, which quantifies the binding of the radioligand to the binding site itself. Here, we quantified expression of the TSPO gene or protein rather than radioligand binding site density. However, we have previously shown that for TSPO, gene expression, protein expression and radioligand binding site data closely correlate^15^. Finally, whilst we present data correlating inducible TSPO expression with the presence of the AP1 binding site in the TSPO core promoter region, to demonstrate causation the AP1 binding site would need to be knocked out from the mouse or rat, and knocked in to a non-*Muroidea* rodent. Furthermore, although we were able to find array expression data for a range of non-rodent mammals that show TSPO is not induced upon myeloid cell activation, we were unable to find array expression data for those rodents that lack the AP1 binding site, such as squirrel or naked mole rat.

In summary, we present *in vitro* expression and sequence alignment data from a range of species, as well as *ex vivo* data from neurodegenerative and neuroinflammatory diseases and associated animal models. We show that inflammation-induced increases in cellular TSPO expression are restricted to microglia from a subset of species within the *Muroidea* superfamily of rodents, and that TSPO is mechanistically linked to classical pro-inflammatory myeloid cell function in mice, but not humans. This challenges the commonly held view that TSPO provides a readout of microglial activation in the human brain and shows that the TSPO PET signal likely reflects the local density of inflammatory cells irrespective of phenotype. The interpretation of TSPO PET data therefore requires revision.

## Methods

### Meta-analysis of TSPO gene expression

Datasets were searched using the search terms “Macrophage/Monocyte/Microglia” and filtered for ‘*Homo sapiens’* and ‘*Mus musculus’*. Datasets with accessible raw data and at least three biological replicates per treatment group were used. To avoid microarray platform-based differences only datasets with Affymetrix chip were used. Raw microarray datasets were downloaded from ArrayExpress (https://www.ebi.ac.uk/arrayexpress/) and RMA normalisation was used. The ‘Limma v.3.42.2’ R package was used to compute differentially expressed genes, and the resulting *P-*values are adjusted for multiple testing with Benjamini and Hochberg’s method to control the false discovery rate^55^. Meta-analysis was performed using R package ‘meta v.5.1.1’. A meta *P*-value was calculated using the random-effect model.

### ChIP-seq data processing and visualisation

ChIP-seq datasets were downloaded from GSE66594^22^ (human) and GSE38377^56^ (mouse). Raw fastq sequences were aligned with Bowtie2 v.2.2.9^57^ to the human reference genome hg19 or to mouse reference genome mm9, annotated SAM files are converted to tag directories using HOMER v.4.11.1^58^ using the makeTagDirectory module. These directories are further used for peak calling using -style histone parameter or converted to the bigWig format normalized to 10^6^ total tag counts with HOMER using the makeUCSCfile module with -fsize parameter set at 2e9. For the analysis of histone ChIP-seq data input samples were utilized as control files during peak detection, whereas IgG control files were used during peak correction of the PU.1 ChIP-seq data. Peaks were visualised using UCSC genome browser^59^.

### Multiple sequence alignment and phylogenetic tree construction

We have retrieved the TSPO promoter region starting from 1 Kbp upstream and 500 bp downstream of the putative transcription start site (TSS) of 34 rodent and non-rodent mammals from ENSEMBL genome database (http://www.ensembl.org/index.htmls). The full list can be found in Supplementary File 2. The multiple sequence alignment was performed using the T-Coffee (v13.45.0.4846264) multiple sequencing tool with the parameter - mode=procoffee which is specifically designed to align the promoter region^60, 61^. The sequence alignment and the phylogenetic tree were visualised using Jalview (v 2.11.1.6)^62^. Phylogenetic tree was constructed using MEGA11 using Maximum Parsimony method with 1000 bootstrap replication. The MP tree was obtained using the Tree-Bisection-Regrafting (TBR) algorithm^63^.

### Motif finding and motif enrichment

We have used SEA (Simple Enrichment Analysis) from the MEME-suite (v 5.4.1) to calculate the relative motif enrichment between Muroidea family species and non-Muroidea mammals^64, 65^. We set the TSPO promoter sequences for the three Muroidea species (Mouse, Rat, Chinese Hamster) as the input sequence and the rest of species as the control sequence. We set the E-value ≤ 10 for calculating significance. We used the motifs for AP1, ETS and SP1 from JASPAR motif database (https://jaspar.genereg.net/).

### Multi-species TSPO expression in macrophage and microglia

Datasets were searched using the search terms “Macrophage/Monocyte”, “Microglia” and “LPS”. Dataset featuring stimulation less than 3 hours were excluded. Datasets with accessible raw data and at least three biological replicates were used. Microarray datasets were analysed as the same way described in section “Meta-analysis of TSPO gene expression”. Raw gene count data for the RNAseq datasets were downloaded from either ArrayExpress or GEO (https://www.ncbi.nlm.nih.gov/geo/) and differential expression was performed using DESeq2 v.1.26.0^66^. For S1a, the mouse *Tspo* expression (GEO ID: GSE38371) fold change was directly used from the respective study since biological replicates were not publicly accessible^23^.

### Human and mouse scRNAseq analysis of microglia

We assessed alterations in gene expression of *TSPO* in human and mouse activated microglia in publicly available scRNAseq datasets. *Postmortem* human brain samples are predominantly studied using single *nucleus* RNA sequencing (snRNAseq) rather than single *cell* RNAseq (sc)RNAseq because the latter requires intact cells which cannot be recovered from frozen brain tissue samples. However, *TSPO* is detected in a very low percentage of nuclei from snRNAseq experiments which prevents accurate assessment of differential expression of *TSPO* across disease or microglial states^54^. For this reason, we searched MEDLINE for human scRNAseq experiments involving AD, MS and ALS donors and mouse brain scRNAseq datasets derived from the respective mouse models, as well as of pro-inflammatory activation with LPS treatment. We found three human studies involving donors with AD^42^ and MS^43, 44^. Where microglia from CSF samples were analysed with scRNAseq. We found no studies with ALS donors. We found three mouse studies: an LPS activated model^39^ an AD model^41^ and acute EAE^40^. A fourth mouse scRNAseq dataset was identified from LPS-treated mice^67^, however, due to its small size (less than 400 microglial cells were sequenced), this dataset was discarded from further analysis. Raw count matrices were downloaded from the Gene Expression Omnibus (GEO) with the following accession numbers: GSE130119^40^, GSE115571^39^, GSE98969^41^, GSE138266^43^ and GSE134578^42^. Data were processed with Seurat (v3)^68^ or nf-core/scflow^69^. Quality control, sample integration, dimension reduction and clustering were performed using default parameters as previously described^54, 70^. Microglial cells (mouse datasets) and microglia-like cells were identified using previously described cell markers. Differential gene expression analysis was performed using MAST^45^ implemented in Seurat to perform zero-inflated regression analysis by fitting a fixed-effects model. Disease vs control group comparisons were performed for all datasets, except for the Keren-Shaul dataset where the AD-associated microglia phenotype was compared to the rest of the microglial population in 5XFAD mice. In all cases, we assessed expression of activated microglial markers. Gene expression alterations were considered significant when the adjusted p value was equal to or lower than 0.05.

### Bulk RNA-seq data preparation and WGCNA network analysis

RAW RNA-seq fastq files for publicly available datasets were downloaded from SRA. Four public human dataset accession are: GSE100382, GSE55536, EMTAB7572, GSE57494 and mouse dataset accession are: GSE103958, GSE62641, GSE82043, GSE58318, E_ERAD_165. The GEO accession ID for the in-house human RNA-seq data is awaiting. Both human and mouse RNA-seq analysis was then performed using nf-core/rnaseq v.1.4.2 pipeline^71^. Human RNA-seq data was aligned to Homo sapiens genome GRCh38 and Mus musculus genome mm10 respectively. Raw count data was first transformed using variance stabilizing transformation (VST) from R package ‘DESeq2 v. 1.26.0’. Genes with an expression value of 1 count in at least 50% of the samples were included in the analysis. Batch correction across datasets were then performed on VST-transformed data using removeBatchEffect function from R package ‘Limma v. 3.42.2’ using the dataset ID as the batch. Batch-corrected normalised data was then used for co-expression network analysis using the R package ‘WGCNA v. 1.69’^72^. The power parameter ranging from 1-20 was screened out using the ‘pickSoftThreshold’ function. A suitable soft threshold of 6 was selected, as it met the degree of independence of 0.85 with the minimum power value. We generated a signed-hybrid network using Pearson correlation with a minimum module size of 30. Subsequently, modules were constructed, and following dynamic branch cutting with a cut height of 0.95. Functional enrichment analysis of the gene modules was performed using the R package ‘WebGestaltR v. 0.4.3’^73^ using default parameters and ‘genome_protein-coding’ as the background geneset.

### Human Brain Tissue

The rapid autopsy regimen of the Netherlands Brain Bank in Amsterdam (coordinator Prof I. Huitinga) was used to acquire the samples. Human tissue was obtained at autopsy from the spinal cord (cervical, thoracic, lumbar levels) from 12 ALS patients, 7 with short disease duration (SDD; <18 months survival; mean survival 11.1 ± 3.4 months) and 4 with medium disease duration (MDD; >24 months survival; mean survival 71.5 ± 31.5 months). Tissues for controls were collected from 10 age-matched cases with no neurological disorders or peripheral inflammation (Table S1). The hippocampal region was collected from 5 AD patients with Braak stage 6, and 5 aged-matched controls that had no cognitive impairments prior to death (Table S2). Active MS lesions were obtained from 5 MS cases as well as white matter from age-matched controls (Table S3). All tissue was collected with the approval of the Medical Ethical Committee of the Amsterdam UMC. All participants or next of kin had given informed consent for autopsy and use of their tissue for research purposes.

## Generation and details of mouse and marmoset models

### Mouse EAE

Spinal cord tissue from mice with EAE was obtained from Biozzi ABH mice housed at Queen Mary University of London, UK (originally obtained from Harlan UK Ltd, Bicester, UK). The mice were raised under pathogen-free conditions and showed a uniform health status throughout the studies. EAE was induced via injection of mouse spinal cord homogenate in complete Freund’s adjuvant (CFA) into mice of 8-12 weeks or 12 months of age as described previously^34, 74^. Immediately, and 24 h after injection mice were given 200ng *Bordetella pertussis* toxin (PT). Age-matched control groups were immunized with CFA and PT. Table S4 gives an overview of the EAE mice used in this study, including a score of neurological signs (0 = normal, 1 = flaccid tail, 2 = impaired righting reflex, 3 = partial hindlimb paresis, 4 = complete hindlimb paresis, 5 = moribund). Spinal cord was collected from acute (aEAE)^74^ in the young mice, and progressive EAE (PEAE) in the 12 month old mice. Animal procedures complied with national and institutional guidelines (UK Animals Scientific Procedures Act 1986) and adhered to the 3R guidelines^75^.

### Marmoset EAE

EAE was induced by subcutaneous immunization with 0.2 g of white matter homogenate emulsified in CFA in 3 adult common marmosets (*Callithrix jacchus*) at 4 dorsal sites adjacent to inguinal and axillary lymph nodes. Animals were monitored daily for clinical symptoms of EAE progression and assigned clinical EAE scores weekly based on extent of disability. Neurological exams were performed by a neurologist prior to each MRI scan. All animals discussed in this study are shown in Table S5. Animal #8 was treated with prednisolone for 5 days as part of a concurrent study (primary results not yet published). These animals were the first within their twin pair that showed three or more brain lesions by *in vivo* MRI and received corticosteroid treatment with the goal to reduce the severity of inflammation and potentially allow longer-term evaluation of the lesions. MRI analyses were performed according to previously published marmoset imaging protocols using T1, T2, T2*, and PD-weighted sequences on a Bruker 7T animal magnet^76^. Marmosets were scanned biweekly over the course of the EAE study. Following the completion of EAE studies, the brains, spinal cords, and optic nerves excised from euthanized animals were scanned by MRI for *postmortem* characterization of brain lesions and previously uncharacterized spinal lesions and optic nerve lesions. Animal procedures complied with national and institutional guidelines (NIH, Bethesda, USA)

### SOD1^G93A^

Female hemizygous transgenic SOD1^G93A^ mice on 129SvHsd genetic background (n=10) and corresponding non transgenic littermates (n=9) were used. This mouse line was raised at the Mario Negri Institute for Pharmacological Research-IRCCS, Milan, Italy, derived from the line (B6SJL-TgSOD1^G93A^-1Gur, originally purchased from Jackson Laboratories, USA) and maintained on a 129S2/SvHsd background^77^. The thoracic segments of spinal cord were collected from 10- and 16-week-old mice and processed as previously described^78^. Briefly, anaesthetised mice were transcardially perfused with 0.1M PBS followed by 4% PFA. The spinal cord was quickly dissected out and left PFA overnight at 4°C, rinsed, and stored 24 h in 10% sucrose with 0.1% sodium azide in 0.1 M PBS at 4°C for cryoprotection, before mounting in optimal cutting temperature compound (OCT) and stored at -80°C.

Procedures involving animals and their care were conducted in conformity with the following laws, regulations, and policies governing the care and use of laboratory animals: Italian Governing Law (D.lgs 26/2014; Authorization 19/2008-A issued 6 March, 2008 by Ministry of Health); Mario Negri Institutional Regulations and Policies providing internal authorization for persons conducting animal experiments; the National Institutes of Health’s Guide for the Care and Use of Laboratory Animals (2011 edition), and European Union directives and guidelines (EEC Council Directive, 2010/63/UE).

### APP^NL-G-F^

For the APP^NL-G-F^ model of AD, male and female brain tissue was obtained from 11 homozygous (APP^NL-G-F/NL-G-F^) APP knock-in mice and 11 wild type mice. Mice were bred at Charles River Laboratories, UK and sampled at the Imperial College London, UK. Brain tissue samples were collected fresh from 10- and 28 week-old mice that were euthanised with sodium pentobarbital and exsanguinated. Animal procedures complied with national and institutional guidelines (UK Animals Scientific Procedures Act 1986) and adhered to 3R guidelines. Hippocampal areas were used as region of interest for characterization.

### Tau^P301S^

Male brain tissue was obtained from 10 homozygous P301S knock-in mice^79–81^ and 8 wild-type C57/Bl6-OLA mice (Envigo, UK) from the Centre for Clinical Brain Sciences, Edinburgh, United Kingdom. Brain tissue samples were collected from 8- and 20-week-old mice that were perfused with PBS and 4% paraformaldehyde, with tissues being post-fixed overnight before being cryopreserved in 30% sucrose and frozen embedded in tissue tec (Leica, UK). Sections were cut, 20μm, on a cryostat onto superfrost plus slides and stored in -80 freezer. Animal procedures complied with national and institutional guidelines (UK Animals Scientific Procedures Act 1986 & University of Edinburgh Animal Care Committees) and adhered to 3R guidelines. Hippocampal areas were used as region of interest for characterization.

For all studies mice were housed 4-5 per standard cages in specific pathogen-free and controlled environmental conditions (temperature: 22±2°C; relative humidity: 55±10% and 12 h of light/dark). Food (standard pellets) and water were supplied *ad libitum*.

### Immunohistochemistry

Paraffin sections were de-paraffinized by immersion in xylene for 5 min and rehydrated in descending concentrations of ethanol and fixed-frozen sections were dried overnight. After washing in PBS, endogenous peroxidase activity was blocked with 0.3 % H_2_O_2_ in PBS while for immunofluorescence sections were incubated in 0.1% glycine. Antigen retrieval was performed with citrate or TRIS/EDTA buffer, depending on the antibody, in a microwave for 3 min at 1000W and 10 min at 180W. Sections were cooled down to RT and incubated with primary antibodies (Table S6) diluted in antibody diluent (Sigma, U3510) overnight. Sections were washed with PBS and afterwards incubated with the appropriate secondary antibodies for 1 h at room temperature. HRP labelled antibodies were developed with diluted 3,3’-diaminobenzidine (DAB; 1:50, DAKO) for 10 min and counterstained with haematoxylin. Sections were immersed in ascending ethanol solutions and xylene for dehydration and mounted with Quick-D. For immunofluorescence, sections were incubated with Alexa Fluor^®^-labelled secondary antibodies. Autofluorescent background signal was reduced by incubating sections in Sudan black (0.1% in 70% EtOH) for 10 min. Nuclei were stained with 4,6-diami-dino-2-phenylindole (DAPI) and slides were mounted onto glass coverslips with Fluoromount^TM^ (Merck).

### Imaging mass cytometry

Antibody conjugation was performed using the Maxpar X8 protocol (Fluidgm). 51 slides of paraffin-embedded tissue from the Medial Temporal Gyrus (MTG) and 48 slides of paraffin-embedded tissue from the Somatosensory Cortex (SSC) underwent IMC staining and ablation. Each slide was within 5-10μm in thickness. The slides underwent routine dewaxing and rehydration before undergoing antigen retrieval, in a pH8 Ethylenediaminetetraacetic acid (EDTA) buffer. The slides were blocked in 10% normal horse serum (Vector Laboratories) before incubation with a conjugated-antibody cocktail (Table S6) at 4⁰C overnight. Slides were then treated in 0.02% Triton X-100 (Sigma-Aldrich) before incubation with an Iridium-intercalator (Fluidigm) then washed in dH2O and air-dried. Image acquisition took place using a Hyperion Tissue Imager (Fluidigm) coupled to a Helios mass cytometer. The instrument was tuned using the manufacturer’s 3-Element Full Coverage Tuning Slide before the slides were loaded into the device. 4 500x500μm regions of interest within the grey matter were selected and then ablated using a laser at a frequency of 200Hz at a 1μm resolution. The data was stored as .mcd files compatible with MCD Viewer software (Fluidigm) then exported as TIFF files. Post-acquisition image processing using ImageJ (v1.53c) software allowed threshold correction and the despeckle function to reduce background noise. The data was opened with HistoCAT (BodenmillerGroup) to quantify the signal of each Ln-channel and exported as .csv files.

### Multiplex immunofluorescence

To immunophenotype microglia/macrophages expressing TSPO in the marmoset CNS, a multi-color multiplex immunofluorescence panel was used to stain for Iba1, PLP, and TSPO. Deparaffinised sections were washed twice in PBS supplemented with 1 mg/ml BSA (PBS/BSA), followed by two washes in distilled water. Antigen retrieval was performed by boiling the slide in 10mM citrate buffer (pH 6) for 10 min in an 800W microwave at maximum power, after which they were allowed to cool for 30 min and washed twice in distilled water. To reduce nonspecific Fc receptor binding, the section was incubated in 250 μl of FcR blocker (Innovex Biosciences, cat. no. NB309) for 15 min at room temperature and washed twice in distilled water. To further reduce background, sections were coated with 250 μl Background Buster (Innovex Biosciences, cat. no. NB306) for 15 min at room temperature and washed twice in distilled water. Sections were incubated for 45 min at room temperature in a primary antibody cocktail containing antibodies diluted in PBS/BSA (Supplemental Table 1), washed in PBS/BSA and three changes of distilled water. They were then incubated for 45 min in a secondary antibody cocktail composed of secondary antibodies diluted in PBS/BSA containing DAPI (Invitrogen, cat. no. D1306, 100 ng/ml) (Supplemental Table 2), then washed once in PBS/BSA and twice in distilled water. To facilitate mounting, the sections were air-dried for 15 min at room temperature, sealed with a coverslip as described previously, and allowed to dry overnight prior to image acquisition.

### Imaging and statistical analyses

Brightfield images were collected at 40x magnification using a Leica DC500 microscope (Leica Microsystems, Heidelberg, Germany, Japan), or a Leica DM6000 (Leica Microsystems, Heidelberg, Germany) or a Zeiss AxioImager.Z2 wide field scanning microscope for fluorescent images. For AD, APP^NL-G-F^, and TAU^P301S^ tissue images were collected from the hippocampus. For ALS tissue, images of the ventral horn and the lateral column were obtained from cervical, thoracic, and lumbar spinal cord levels. For mouse EAE and SOD1^G93A^ mice, images of grey and white matter of the spinal cord were collected per case. ImageJ software was used for picture analyses. Nuclei and stained cells were counted manually using the cell counter plugin (de Vos, University of Sheffield, UK), excluding nuclei at the rim of each picture and within blood vessels. To determine inter-observer variation 18 pictures were manually counted by 3 independent observers with a correlation coefficient of > 0.9. To determine single cell TSPO expression, IBA+ or GFAP+ cells were outlined manually using the imageJ using the ROI manager. Afterwards TSPO+ pixels were measured within IBA+ and GFAP+ ROIs per cell. Data were analyzed using GraphPad Prism 9.1.0 software. All data were tested for normal distribution, using the Shapiro-Wilk normality test. Significant differences were detected using an unpaired t-test or one-way analysis of variance test. Dunnett’s post-hoc test was performed to analyze which groups differ significantly. Number of mice were calculated by power analysis and as a maximum 6-8 mice were used per group based on previous studies^34^. Data was considered significant when P < 0.05.

### BV2 and primary mouse macrophage culture

All cells were kept at 37°C, 5% CO_2_ and 95% humidity. Mouse BV2 cells (a kind gift from Federico Roncaroli, Manchester) were cultured in RPMI-1640 containing 2mM GlutaMAX and 10% heat inactivated FBS (all Gibco). For experiments BV2 were seeded at 1x10^4 cells per well of a 96-well plate the day before treatment. Primary mouse bone marrow-derived macrophages (BMDMs) were obtained from bone marrow of adult C57BL/6 mice and cultured in DMEM containing 10% FBS, penicillin/streptomycin, and glutamine supplemented with M-CSF (10ng/mL; Peprotech) as previously described (Ying et al. 2013). All animal procedures were approved by the Memorial University Animal Care Committee in accordance with the guidelines set by the Canadian Council in Animal Care.

### Primary human macrophage culture

All donors gave informed consent under a REC approved protocol (12/LO/0538). Human monocyte derived macrophages (MDMs) were obtained from fresh blood of male and female, healthy donors between 20 and 60 years after CD14-affinity purification. In brief, whole blood was diluted 1:1 with DPBS (Sigma), layered onto Ficoll (Sigma) and spun for 20 min at 800xg with minimal acceleration/deceleration. Peripheral mononuclear cells were collected, washed, and labelled with CD14-affinity beads (Miltenyi) according to the manufacturers protocol. CD14 monocytes were eluted and cultured at 5x10^5 cells/ml in RPMI-1640 containing 2mM GlutaMAX, 10% heat inactivated FBS, and 25ng/ml M-CSF (all Gibco) with medium change after 3 days. MDMs were used after 7 days in-vitro culture. For monocytes, M-CSF was omitted from the medium and cells were used immediately ex-vivo.

### Human TSPO genotyping

Genotyping at rs6971 was performed by LGC. Where not specified, studies were performed with homozygous A carriers due to the high affinity for XBD-173 (high-affinity binders; HAB). Homozygous T carriers were grouped as low affinity binders (LAB). Heterozygous rs6971 carriers were omitted from this study.

### iPSC culture and microglia-like cell differentiation

The human induced pluripotent stem cell (iPSC) line SFC841-03-01 (https://hpscreg.eu/cell-line/STBCi044-A, previously derived from a healthy donor^82^, Oxford Parkinson’s Disease Centre/StemBANCC) was obtained under MTA from the James Martin Stem Cell Facility, University of Oxford and cultured in feeder-free, fully defined conditions. In brief, iPSCs were maintained in E8 medium on Geltrex (both Gibco) and fed every day until 80% confluent. For cell cluster propagation, iPSCs were lifted with 0.5 mM EDTA (Thermo) in DPBS and upon visible dissociation, EDTA was removed, and iPSC were diluted 4-6 times in E8 for culture maintenance. iPSCs were screened genotypically for chromosomal abnormalities using single nucleotide polymorphism analysis and phenotypically using Nanog (Cell Signalling) and Tra-1-60 (BioLegend) immune positivity. Mycoplasma infection was excluded based on LookOut test (Sigma) according to manufacturer’s protocol. Microglia-like cells were differentiated according to Haenseler et al 2017^83^. In short, on day 0 iPSCs were dissociated with TrypLE Express (Gibco) and 4x10^6 iPSCs were added to one well of 24-well AggreWell^TM^ 800 (Stem Cell Technology) according to the manufacturer’s protocol in 2ml EB medium (E8, SCF (20ng/ml, Miltenyi), BMP4 (50ng/ml; Gibco), VEGF (50ng/ml, PeproTech)) with 10uM ROCK inhibitor (Y-27632, Abcam). From day 1 to 6, 75% medium was exchanged with fresh EB. On day 7 embryoid bodies were transferred to 2x T175 flasks containing factory medium (XVIVO-15 (Lonza), 2mM GlutaMAX, 50uM 2-Mercaptoethanol, 25ng/ml IL-3, and 100ng/ml M-CSF (all Gibco)) and fed weekly with factory medium. Starting from week 4 after transfer, medium was removed and tested for the presence of primitive macrophages using CD45 (immunotools), CD14 (immunotools) and CD11b (Biolegend) immunopositivity by flow cytometry (FACSCalibur, BD Biosciences). Primitive macrophages were transferred to microglia medium (SILAC Adv DMEM/F12 (Gibco), 10 mM glucose (Sigma), 2 mM GlutaMAX, 0.5 mM L-lysine (Sigma), 0.5 mM L-arginine (Sigma), 0.00075% phenol red (Sigma), 100ng/ml IL-34 (PeproTech), 10gn/ml GM-CSF (Gibco)), fed every 3-4 days and used for experiments after 7 days.

### Drug treatments and Cell activation

Cells were treated with XBD-173 at the indicated concentrations for 1h prior to LPS activation or for 20h prior to phagocytosis. Pro-inflammatory activation was induced with lipopolysaccharide (100ng/ml; Sigma) for 24h. For live-cell phagocytosis assays, pHrodo®-labelled zymosan A bioparticles (Thermo) were added to the culture medium and incubated for 2h at 37°C with 5% CO_2_. pHrodo®-fluorescence intensity was acquired in a plate reader (Cytation5, BioTek) or by Flow cytometry (FACSCalibur, BD Biosciences).

### Cytokine analysis

Cytokines were assessed from cell-free cell culture supernatant using enzyme-linked immunosorbent assay (ELISA) according to the manufacturers’ protocols. The following assays were used: mouse-TNFα and mouse-IL-6 ELISA (R&D Systems), huma-TNFα and human-IL-6 (BD Biosciences). Absorbance was measured in a Spark plate reader (Tecan).

### RNA Sequencing\

RNA was extracted from control and LPS treated (100ng/mL, 24 hours) primary human macrophages using the RNeasy Mini Kit. cDNA libraries (Total RNA with rRNA depletion) were prepared and sequenced using a HiSeq4000. Lanes were run as 75 bases Paired End. Sequencing depth was minimum 40 million reads per sample

### LC-MSMS analysis of supernatant for XBD173 concentration

Supernatant samples were stored at -20°C or lower until analysis. Samples (25 µL) were prepared for analysis by protein precipitation with acetonitrile containing internal standard (tolbutamide) (200 µL) followed by mixing (150 rpm, 15 min) and centrifugation (3000 rpm, 15 min). The supernatant (50) µL was diluted with water (100 µL) and mixed (100 rpm, 15min). Samples were analysed by LC-MSMS (Shimadzu Nexera X2 UHPLC/Shimadzu LCMS 8060) with Phenomenex Kinetex Biphenyl (50 x 2.1)mm, 1.7 µm column and mobile phase components water/0.1% formic acid (A) and acetonitrile/0.1% formic acid (B). Mobile phase gradient was 0 to 0.3 min 2% B; 0.3 to 1.1 min increase to 95% B; 1.1 to 1.75 min 95% B, 1.75 to 1.8 min decrease to 2% B; 1.8 to 2.5 min 2% B. Flow rate was 0.4 mL/min. Injection volume was 1 µL. Calibration standards were prepared by spiking XDB173 into control supernatant over the range 2-10000 ng/mL, then preparing and analysing as for the study samples. Lower limit of detection was 2 ng/mL.

## Acknowledgements

The authors thank the UK MS society for financial support (grant number: C008-16.1). Raw count matrices corresponding to microglia-like cells from Ramesh et al.,^44^ were kindly provided by Prof Michael Wilson. DRO was funded by an MRC Clinician Scientist Award (MR/N008219/1). PMM acknowledges generous support from Edmond J Safra Foundation and Lily Safra, the NIHR Investigator programme and the UK Dementia Research Institute. PMM and DRJO thank the Imperial College Healthcare Trust-NIHR Biomedical Research Centre for infrastructure support and the Medical Research Council for support of TSPO studies. Dr Sally Cowley (Oxford Parkinson’s Disease Centre, James Martin Stem Cell Facility, University of Oxford) provided the iPS cell line and expertise in differentiation to iPS-microglia.

## Reporting Summary

Further information on research design is available in the Nature Research Reporting Summary linked to this article.

## Data availability

The data that support the findings of this study are available in this manuscript and the Supplementary Information. Source data are provided with this paper.

## Code availability

Code used throughout this study is available upon request from the corresponding authors.

## Author contributions

Conceptualisation: E.N., N.F., M.W., S.A., and D.R.O. Technical and Analysis Support: J.A., D.S., S.C., M.C.T., T.Saito., T.Saido., M.W., C.S.M., C.B., and C.I.R. Data Collection and Curation: E.N., N.F., M.W., M.C.M., S.T., R.C.J.M., I.F., J.B., D.H., and R.P. Writing – Original Draft: E.N., N.F., S.A., and D.R.O. Writing – Review and Editing: All authors have reviewed the manuscript. Visualisation: E.N., N.F., M.W., M.C.M., S.T., R.C.J.M., and I.F. Supervision: S.A., and D.R.O.

---

**Figure S1.**
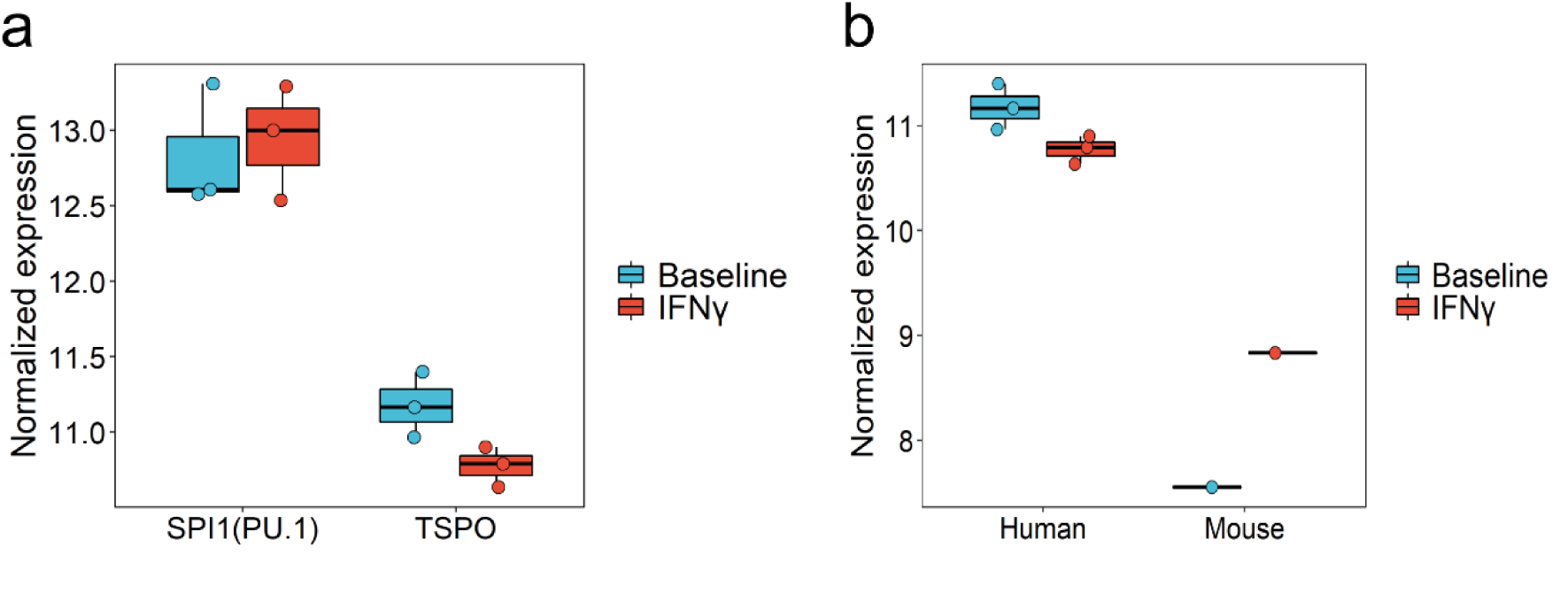
**a** Boxplot showing TSPO fold change in human and mouse macrophages in baseline and IFNγ treated samples. **b** Boxplot showing PU.1 (SPI1) transcription factor and TSPO gene expression change in IFNγ treated macrophage compared to baseline condition.

**Figure S2.**
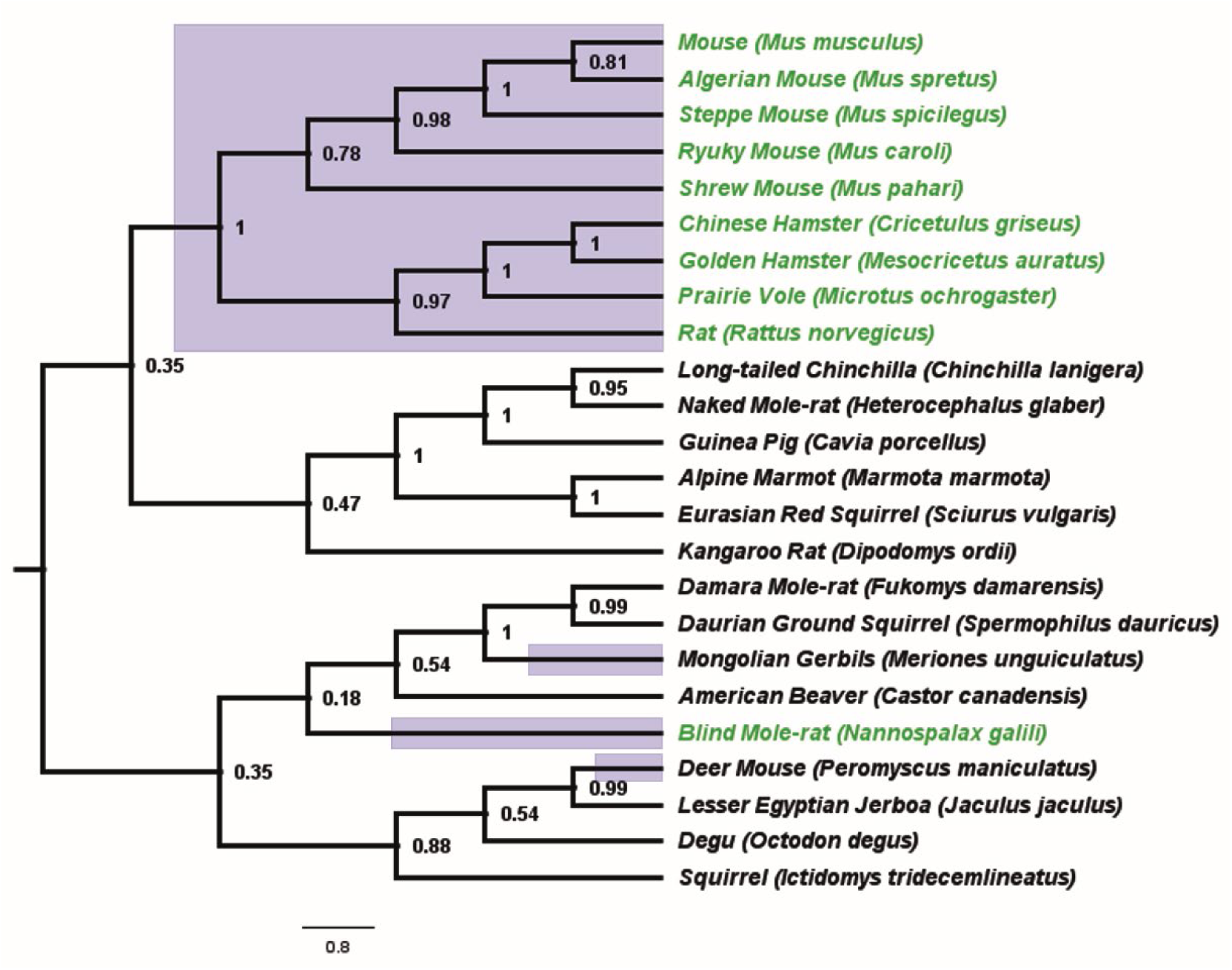
Of the 24 rodent species examined here, 12/24 are from the Muroidea superfamily (purple branches). 10 of these 12 Muroidea species contain the AP1 binding site in the TSPO promoter (Green Highlight). We did not find any rodent species outside the Muroidea superfamily that contain the AP1 binding site in the TSPO promoter. The phylogenetic analysis shows that majority of the species (9/12) from Muroidea superfamily forms a single clade. Phylogenetic tree was generated using the Maximum Parsimony method in MEGA11. The consistency index (CI) is 0.623399 (0.553120) and the retention index (RI) is 0.525671 (0.525671) for all sites and parsimony-informative sites (in parentheses). The percentage of replicate trees in which the associated taxa clustered together in the bootstrap test (1000 replicates) are shown next to the branches.

**Figure S3.**
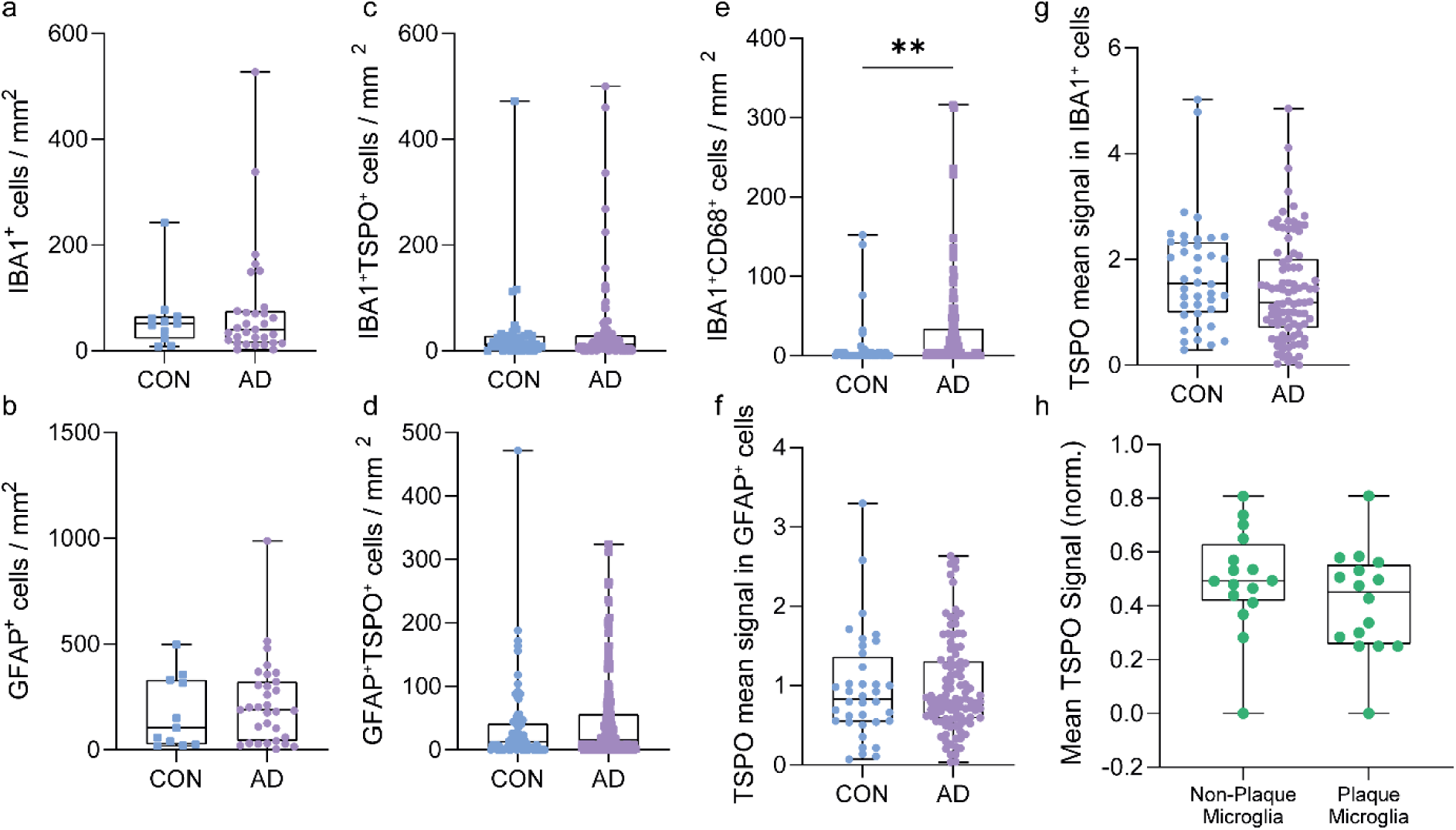
**a-d** no increase in total or TSPO+ microglia (P) and astrocytes (P) are observed in control versus AD. **e** An increase in CD68+IBA1+ cells is observed in AD. **f,g** No increases in mean TSPO signal in microglia and astrocytes is observed in AD relative to control. **h** No differences are observed in mean TSPO signal in microglia associated with plaques compared to mean TSPO signal in microglia that are distant from plaques.

**Figure S4.**
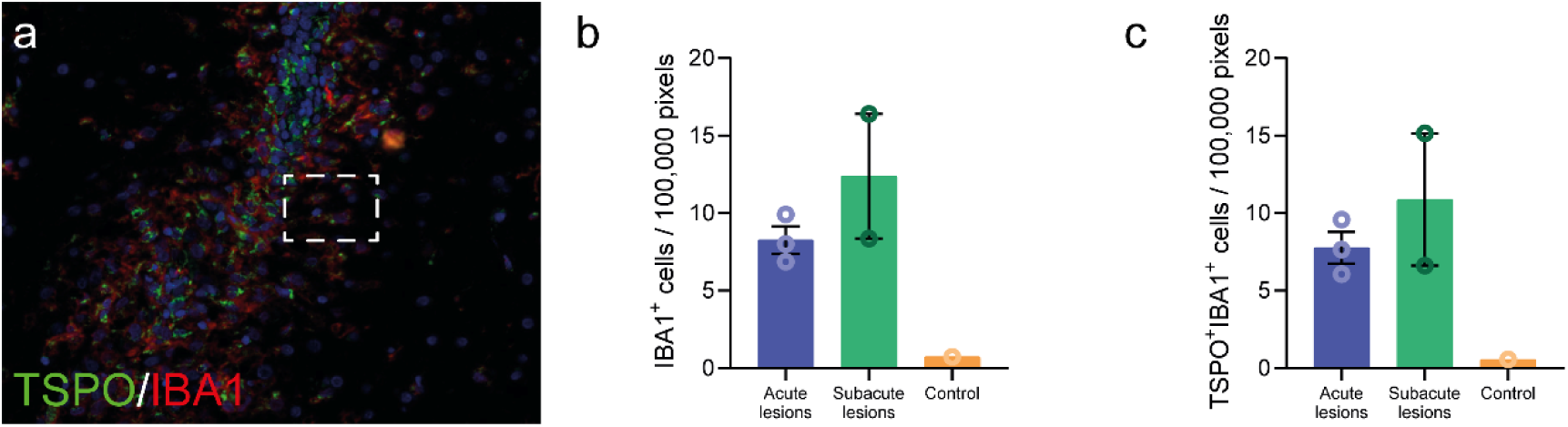
**a** Representative image of an acute lesion in marmoset EAE. IBA1+ and TSPO+IBA1+ cells are increased in acute and subacute lesions compared to white matter in control marmoset.

**Figure S5.**
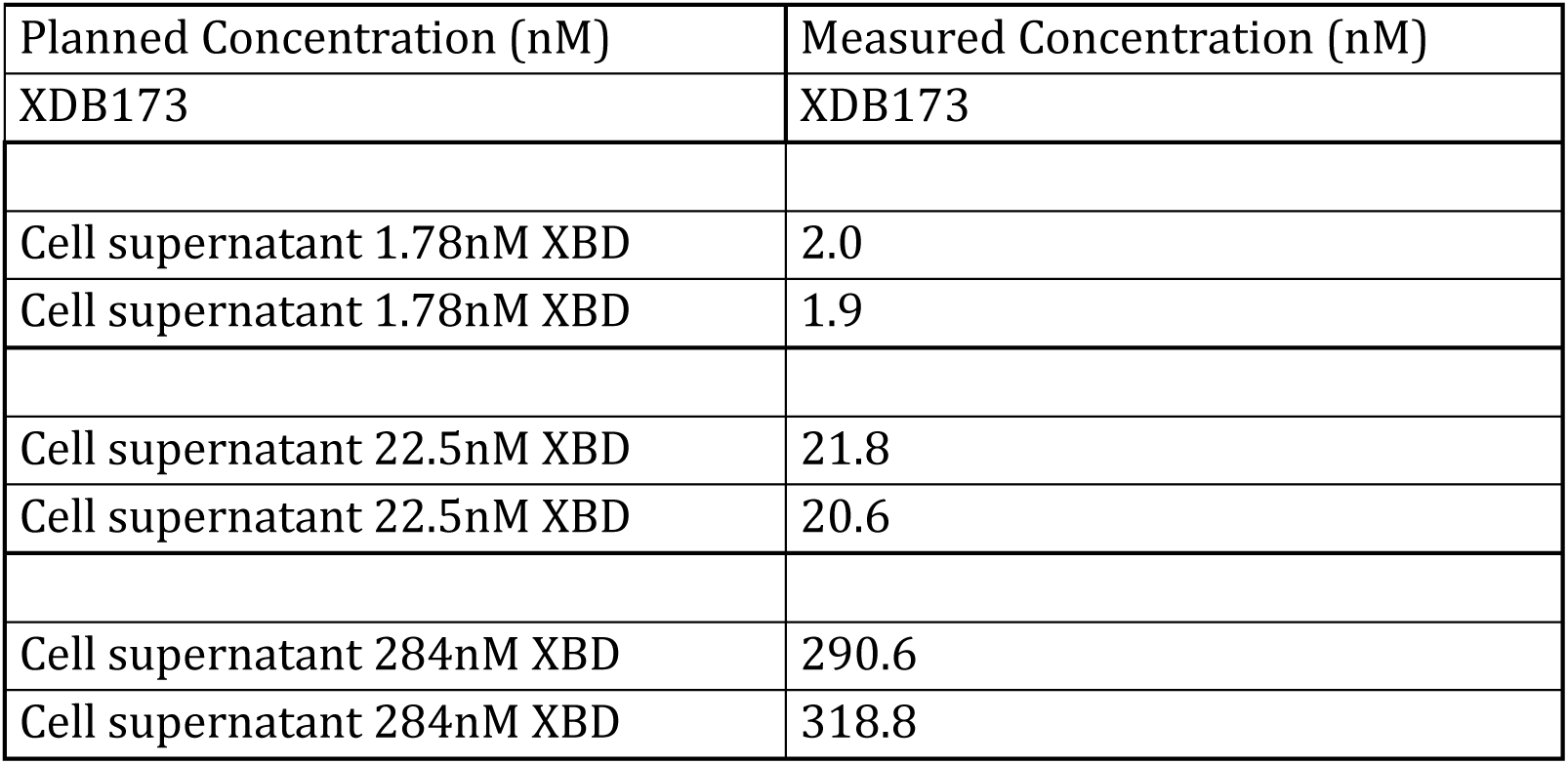
Planned and measured concentrations of XBD173 in medium for experiments described in Figures 8d and 8e.

**Supplementary Table 1.** Clinical details of AD and control cases

| Case | Age/sex | Diagnosis | Region | Braak stage | PMD, h:min |
| --- | --- | --- | --- | --- | --- |
| AD cases |  |  |  |  |  |
| 1 | 81/F | AD | HC (anterior) | 6 | 05:30 |
| 2 | 88/F | AD | HC (anterior) | 6 | 06:19 |
| 3 | 62/M | AD | HC (anterior) | 6 | 06:15 |
| 4 | 64/F | AD | HC (anterior) | 6 | 06:30 |
| 5 | 76/M | AD | HC (anterior) | 6 | 04:40 |
| Controls |  |  |  |  |  |
| 1 | 65/F | NDC | HC | 2 | 07:10 |
| 2 | 90/F | NDC | HC | 3 | 06:10 |
| 3 | 81/F | NDC | HC | 3 | 05:30 |
| 4 | 77/M | NDC | HC (anterior) | 2 | 04:30 |
| 5 | 81/F | Ischemic changes | HC (anterior) | 4 | 05:50 |

Abbreviations: F – female; HC – hippocampus; M – male; NDC – non-demented control; PMD – postmortem delay.

**Supplementary Table 2.**
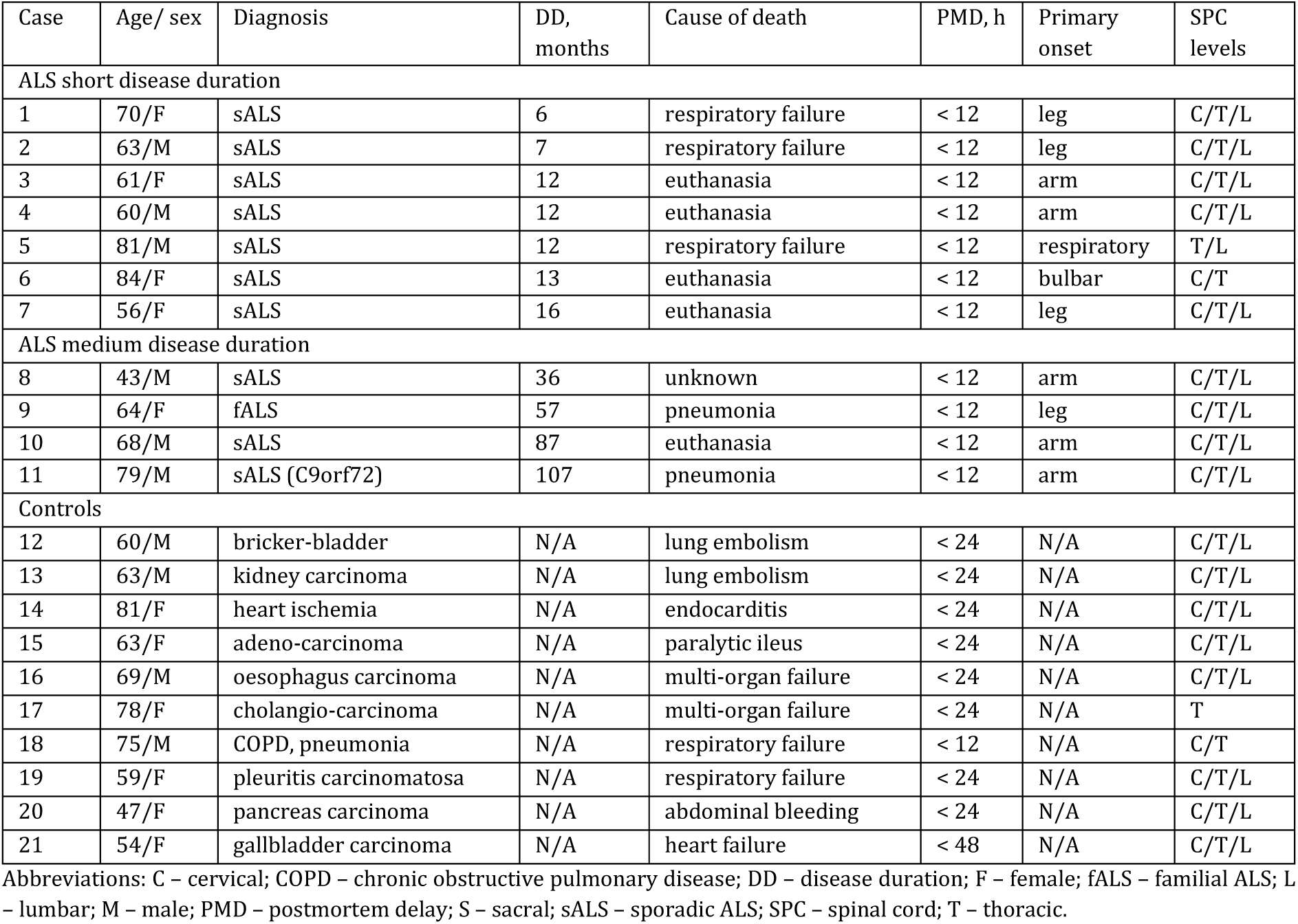
Clinical details of ALS and control cases

| Case | Age/ sex | Diagnosis | DD, months | Cause of death | PMD, h | Primary onset | SPC levels |
| --- | --- | --- | --- | --- | --- | --- | --- |
| ALS short disease duration |  |  |  |  |  |  |  |
| 1 | 70/F | sALS | 6 | respiratory failure | < 12 | leg | C/T/L |
| 2 | 63/M | sALS | 7 | respiratory failure | < 12 | leg | C/T/L |
| 3 | 61/F | sALS | 12 | euthanasia | < 12 | arm | C/T/L |
| 4 | 60/M | sALS | 12 | euthanasia | < 12 | arm | C/T/L |
| 5 | 81/M | sALS | 12 | respiratory failure | < 12 | respiratory | T/L |
| 6 | 84/F | sALS | 13 | euthanasia | < 12 | bulbar | C/T |
| 7 | 56/F | sALS | 16 | euthanasia | < 12 | leg | C/T/L |
| ALS medium disease duration |  |  |  |  |  |  |  |
| 8 | 43/M | sALS | 36 | unknown | < 12 | arm | C/T/L |
| 9 | 64/F | fALS | 57 | pneumonia | < 12 | leg | C/T/L |
| 10 | 68/M | sALS | 87 | euthanasia | < 12 | arm | C/T/L |
| 11 | 79/M | sALS (C9orf72) | 107 | pneumonia | < 12 | arm | C/T/L |
| Controls |  |  |  |  |  |  |  |
| 12 | 60/M | bricker-bladder | N/A | lung embolism | < 24 | N/A | C/T/L |
| 13 | 63/M | kidney carcinoma | N/A | lung embolism | < 24 | N/A | C/T/L |
| 14 | 81/F | heart ischemia | N/A | endocarditis | < 24 | N/A | C/T/L |
| 15 | 63/F | adeno-carcinoma | N/A | paralytic ileus | < 24 | N/A | C/T/L |
| 16 | 69/M | oesophagus carcinoma | N/A | multi-organ failure | < 24 | N/A | C/T/L |
| 17 | 78/F | cholangio-carcinoma | N/A | multi-organ failure | < 24 | N/A | T |
| 18 | 75/M | COPD, pneumonia | N/A | respiratory failure | < 12 | N/A | C/T |
| 19 | 59/F | pleuritis carcinomatosa | N/A | respiratory failure | < 24 | N/A | C/T/L |
| 20 | 47/F | pancreas carcinoma | N/A | abdominal bleeding | < 24 | N/A | C/T/L |
| 21 | 54/F | gallbladder carcinoma | N/A | heart failure | < 48 | N/A | C/T/L |

Abbreviations: C – cervical; COPD – chronic obstructive pulmonary disease; DD – disease duration; F – female; fALS – familial ALS; L – lumbar; M – male; PMD – postmortem delay; S – sacral; sALS – sporadic ALS; SPC – spinal cord; T – thoracic.

**Supplementary Table 3.** Clinical details of MS and control cases

| Case | Age/sex | Diagnosis | Disease duration, years | Cause of death | PMD, h:min |
| --- | --- | --- | --- | --- | --- |
| MS cases |  |  |  |  |  |
| 1 | 35/F | SPMS | 10 | Euthanasia | 10:20 |
| 2 | 54/F | SPMS | 27 | Respiratory failure | 9:25 |
| 3 | 50/F | SPMS | 18 | Euthanasia | 9:05 |
| 4 | 50/M | SPMS | 21 | Unknown | 10:50 |
| 5 | 63/F | Unknown | Unknown | Unknown | 10:50 |
| Controls |  |  |  |  |  |
| 1 | 84/M | NNC | N/A | Heart failure | 5:35 |
| 2 | 89/F | NNC | N/A | Pneumonia | 3:52 |
| 3 | 79/M | NNC | N/A | Heart failure | 6:20 |
| 4 | 73/F | NNC | N/A | Mamma carcinoma | 7:45 |
| 5 | 87/F | NNC | N/A | Pneumonia | 7:00 |

Abbreviations: F – female; M – male; N/A – not applicable; NNC – non-neurological control; PMD – postmortem delay; SPMS – secondary progressive multiple sclerosis.

**Supplementary Table 4.**
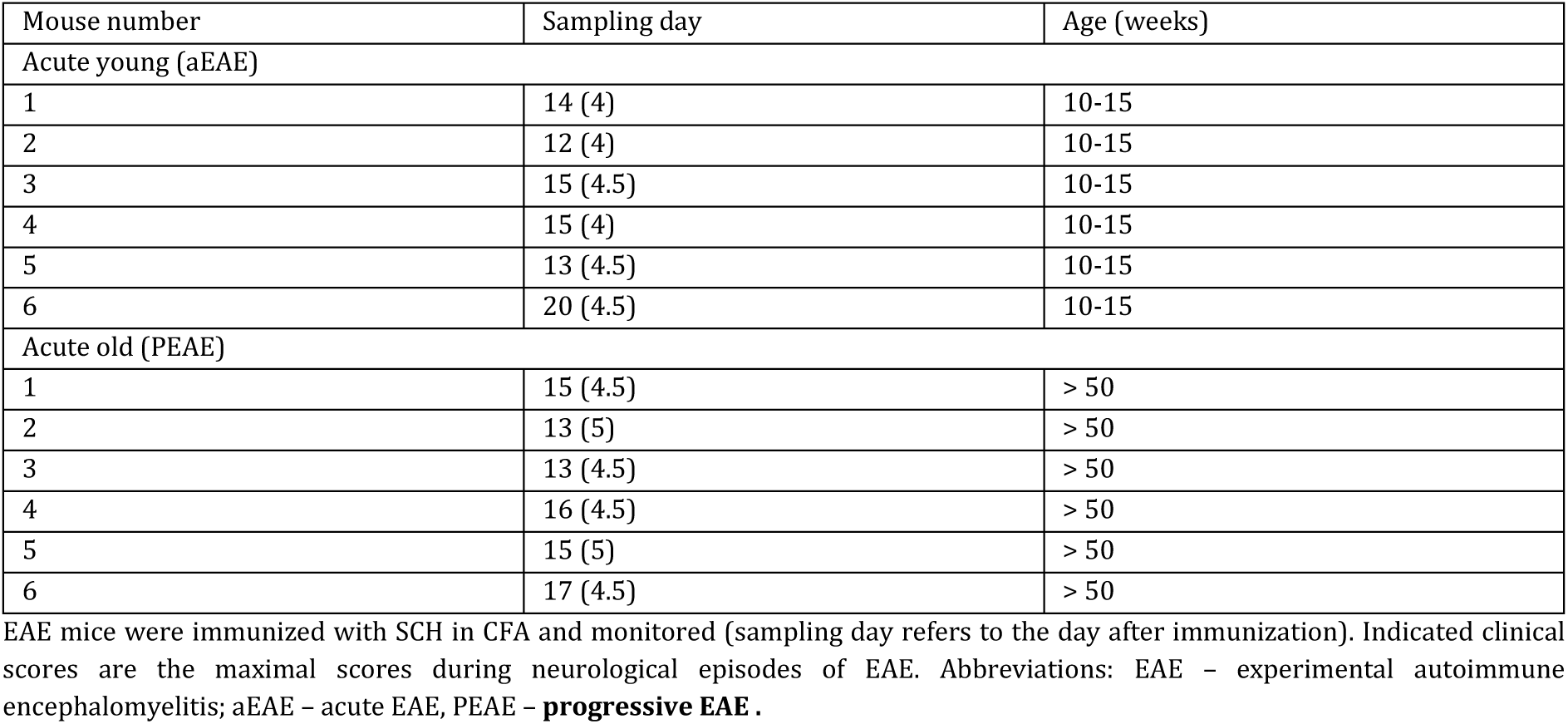
Clinical History of mice with EAE

| Mouse number | Sampling day | Age (weeks) |
| --- | --- | --- |
| Acute young (aEAE) |  |  |
| 1 | 14 (4) | 10-15 |
| 2 | 12 (4) | 10-15 |
| 3 | 15 (4.5) | 10-15 |
| 4 | 15 (4) | 10-15 |
| 5 | 13 (4.5) | 10-15 |
| 6 | 20 (4.5) | 10-15 |
| Acute old (PEAE) |  |  |
| 1 | 15 (4.5) | > 50 |
| 2 | 13 (5) | > 50 |
| 3 | 13 (4.5) | > 50 |
| 4 | 16 (4.5) | > 50 |
| 5 | 15 (5) | > 50 |
| 6 | 17 (4.5) | > 50 |

EAE mice were immunized with SCH in CFA and monitored (sampling day refers to the day after immunization). Indicated clinical scores are the maximal scores during neurological episodes of EAE. Abbreviations: EAE – experimental autoimmune encephalomyelitis; aEAE – acute EAE, PEAE – **progressive EAE**.

**Supplementary Table 5.** Clinical History of Marmosets

| Animal ID | Gender | Disease Status | Age at EAE induction (years) | Disease duration (days) | Age (years) |
| --- | --- | --- | --- | --- | --- |
| 1 | M | Control | N/A | N/A | 3 |
| 4 | F | EAE | 2.0 | 32 | 2.1 |
| 5 | F | EAE | 1.6 | 105 | 1.9 |
| 8 | F | EAE | 1.6 | 123 | 1.9 |

Abbreviations: EAE – experimental autoimmune encephalitis; N/A – not applicable.

**Supplementary Table 6.** Antibodies for immunohistochemistry and imaging mass cytometry

| Antigen | Species (isotype) | Clonality | Dilution | Antigen Retrieval | Product Number | Supplier |
| --- | --- | --- | --- | --- | --- | --- |
| TSPO | goat | pAb | 1:750 | Citrate | NB100-41398 | Novus Biologicals |
| TSPO | rabbit | mAb | 1:750 | Citrate | AB109497 | Abcam |
| IBA1 | rabbit | pAb | 1:10000 | Tris-EDTA | 019-19741 | Wako |
| IBA1 | goat | pAb | 1:1000 | Tris-EDTA | AB48004 | Abcam |
| IBA1 | guinea pig | pAb | 1:100 | Citrate | 234004 | Synaptic Systems |
| GFAP | chicken | pAb | 1:500 | Citrate | AB5541 | Millipore |
| HLA-DR | mouse (IgG2B) | mAb | 1:750 | Citrate | 14-9956-82 | Invitrogen |
| Aβ IC16 | mouse (IgG2A) | mAb | 1:400 | Citrate | N/A | in-house<sup>a</sup> |
| P-Tau AT8 | mouse (IgG1) | mAb | 1:400 | Citrate | AB_223647 | Invitrogen |
| PLP | mouse (IgG2A) | mAb | 1:200 | Citrate | MCA839G | Bio-Rad |
| IMC | Ln-Isotope |  |  |  |  |  |
| CD68 | 159Tb |  | 1:800 | EDTA | 3159035D | Fluidigm |
| GFAP | 162Dy |  | 1:600 | EDTA | Ab218309 | Abcam |
| HLA-DR | 174Yb |  | 1:400 | EDTA | 3174025D | Fluidigm |
| IBA1 | 169Tm |  | 1:3000 | EDTA | 019-197471 | Wako |
| TSPO | 149Sm |  | 1:400 | EDTA | Ab213654 | Abcam |

<sup>a</sup>With permission from Carsten Korth, Heinrich Heine University, Düsseldorf, Germany. Abbreviations: GFAP – glial fibrillary acidic protein; IBA1 – ionized calcium-binding adaptor molecule 1; mAb – monoclonal antibody; pAb – polyclonal antibody; P-Tau – phosphorylated Tau (Ser202, Thr205).

## References

1. Cunningham, C. Microglia and neurodegeneration: the role of systemic inflammation. Glia 61, 71–90 (2013).

2. Heneka, M.T., et al. Locus ceruleus controls Alzheimer’s disease pathology by modulating microglial functions through norepinephrine. Proc Natl Acad Sci U S A 107, 6058–6063 (2010).

3. O’Sullivan, J.B., Ryan, K.M., Curtin, N.M., Harkin, A. & Connor, T.J. Noradrenaline reuptake inhibitors limit neuroinflammation in rat cortex following a systemic inflammatory challenge: implications for depression and neurodegeneration. Int J Neuropsychopharmacol 12, 687–699 (2009).

4. Brown, G.C. & Vilalta, A. How microglia kill neurons. Brain Res 1628, 288–297 (2015).

5. Brown, G.C. & Neher, J.J. Inflammatory neurodegeneration and mechanisms of microglial killing of neurons. Mol Neurobiol 41, 242–247 (2010).

6. Brown, G.C. & Neher, J.J. Microglial phagocytosis of live neurons. Nature reviews. Neuroscience 15, 209–216 (2014).

7. Guilarte, T.R. TSPO in diverse CNS pathologies and psychiatric disease: A critical review and a way forward. Pharmacol Ther 194, 44–58 (2019).

8. Guilarte, T.R., Rodichkin, A.N., McGlothan, J.L., Acanda De La Rocha, A.M. & Azzam, D.J. Imaging neuroinflammation with TSPO: A new perspective on the cellular sources and subcellular localization. Pharmacol Ther, 108048 (2021).

9. Pascoal, T.A., et al. Microglial activation and tau propagate jointly across Braak stages. Nature medicine 27, 1592–1599 (2021).

10. Jucaite, A., et al. Effect of the myeloperoxidase inhibitor AZD3241 on microglia: a PET study in Parkinson’s disease. Brain : a journal of neurology 138, 2687–2700 (2015).

11. Bae, K.R., Shim, H.J., Balu, D., Kim, S.R. & Yu, S.W. Translocator protein 18 kDa negatively regulates inflammation in microglia. J Neuroimmune Pharmacol 9, 424–437 (2014).

12. Wang, M., et al. Macroglia-microglia interactions via TSPO signaling regulates microglial activation in the mouse retina. J Neurosci 34, 3793–3806 (2014).

13. Karlstetter, M., et al. Translocator protein (18 kDa) (TSPO) is expressed in reactive retinal microglia and modulates microglial inflammation and phagocytosis. J Neuroinflammation 11, 3 (2014).

14. Gottfried-Blackmore, A., Sierra, A., Jellinck, P.H., McEwen, B.S. & Bulloch, K. Brain microglia express steroid-converting enzymes in the mouse. J Steroid Biochem Mol Biol 109, 96–107 (2008).

15. Owen, D.R., et al. Pro-inflammatory activation of primary microglia and macrophages increases 18 kDa translocator protein expression in rodents but not humans. J Cereb Blood Flow Metab 37, 2679–2690 (2017).

16. Nutma, E., et al. Activated microglia do not increase 18 kDa translocator protein (TSPO) expression in the multiple sclerosis brain. Glia 69, 2447–2458 (2021).

17. Srivastava, P.K., Hull, R.P., Behmoaras, J., Petretto, E. & Aitman, T.J. JunD/AP1 regulatory network analysis during macrophage activation in a rat model of crescentic glomerulonephritis. BMC Syst Biol 7, 93 (2013).

18. Saito, T., et al. Single App knock-in mouse models of Alzheimer’s disease. Nature neuroscience 17, 661–663 (2014).

19. Yoshiyama, Y., et al. Synapse loss and microglial activation precede tangles in a P301S tauopathy mouse model. Neuron 53, 337–351 (2007).

20. Gurney, M.E., et al. Motor neuron degeneration in mice that express a human Cu,Zn superoxide dismutase mutation. Science 264, 1772–1775 (1994).

21. Baker, D., et al. Induction of chronic relapsing experimental allergic encephalomyelitis in Biozzi mice. J Neuroimmunol 28, 261–270 (1990).

22. Schmidt, S.V., et al. The transcriptional regulator network of human inflammatory macrophages is defined by open chromatin. Cell Res 26, 151–170 (2016).

23. Ostuni, R., et al. Latent enhancers activated by stimulation in differentiated cells. Cell 152, 157–171 (2013).

24. Shlyueva, D., Stampfel, G. & Stark, A. Transcriptional enhancers: from properties to genome-wide predictions. Nat Rev Genet 15, 272–286 (2014).

25. Celada, A., et al. The transcription factor PU.1 is involved in macrophage proliferation. J Exp Med 184, 61–69 (1996).

26. Ghisletti, S., et al. Identification and characterization of enhancers controlling the inflammatory gene expression program in macrophages. Immunity 32, 317–328 (2010).

27. Rashid, K., Geissl, L., Wolf, A., Karlstetter, M. & Langmann, T. Transcriptional regulation of Translocator protein (18kDa) (TSPO) in microglia requires Pu.1, Ap1 and Sp factors. Biochim Biophys Acta Gene Regul Mech 1861, 1119–1133 (2018).

28. Lane, C.A., Hardy, J. & Schott, J.M. Alzheimer’s disease. Eur J Neurol **25**, 59–70 (2018).

29. Tiwari, S., Atluri, V., Kaushik, A., Yndart, A. & Nair, M. Alzheimer’s disease: pathogenesis, diagnostics, and therapeutics. International journal of nanomedicine 14, 5541–5554 (2019).

30. Kellner, A., et al. Autoantibodies against beta-amyloid are common in Alzheimer’s disease and help control plaque burden. Ann Neurol 65, 24–31 (2009).

31. Hansen, D.V., Hanson, J.E. & Sheng, M. Microglia in Alzheimer’s disease. J Cell Biol 217, 459–472 (2018).

32. Xuan, F.L., Chithanathan, K., Lillevali, K., Yuan, X. & Tian, L. Differences of Microglia in the Brain and the Spinal Cord. Front Cell Neurosci 13, 504 (2019).

33. Nutma, E., et al. A quantitative neuropathological assessment of translocator protein expression in multiple sclerosis. Brain : a journal of neurology 142, 3440–3455 (2019).

34. Peferoen, L.A., et al. Ageing and recurrent episodes of neuroinflammation promote progressive experimental autoimmune encephalomyelitis in Biozzi ABH mice. Immunology 149, 146–156 (2016).

35. Tuisku, J., et al. Effects of age, BMI and sex on the glial cell marker TSPO - a multicentre [(11)C]PBR28 HRRT PET study. European journal of nuclear medicine and molecular imaging 46, 2329–2338 (2019).

36. Gaitan, M.I., et al. Perivenular brain lesions in a primate multiple sclerosis model at 7-tesla magnetic resonance imaging. Mult Scler 20, 64–71 (2014).

37. . t Hart, B.A., Vogels, J., Bauer, J., Brok, H.P. & Blezer, E. Non-invasive measurement of brain damage in a primate model of multiple sclerosis. Trends Mol Med 10, 85–91 (2004).

38. Lefeuvre, J.A., et al. The spectrum of spinal cord lesions in a primate model of multiple sclerosis. Mult Scler 26, 284–293 (2020).

39. Sousa, C., et al. Single-cell transcriptomics reveals distinct inflammation-induced microglia signatures. EMBO Rep 19, e46171 (2018).

40. Wheeler, M.A., et al. MAFG-driven astrocytes promote CNS inflammation. Nature 578, 593–599 (2020).

41. Keren-Shaul, H., et al. A Unique Microglia Type Associated with Restricting Development of Alzheimer’s Disease. Cell 169, 1276–1290 e1217 (2017).

42. Gate, D., et al. Clonally expanded CD8 T cells patrol the cerebrospinal fluid in Alzheimer’s disease. Nature 577, 399–404 (2020).

43. Schafflick, D., et al. Integrated single cell analysis of blood and cerebrospinal fluid leukocytes in multiple sclerosis. Nat Commun 11, 247 (2020).

44. Ramesh, A., et al. A pathogenic and clonally expanded B cell transcriptome in active multiple sclerosis. Proc Natl Acad Sci U S A 117, 22932–22943 (2020).

45. Finak, G., et al. MAST: a flexible statistical framework for assessing transcriptional changes and characterizing heterogeneity in single-cell RNA sequencing data. Genome Biol 16, 278 (2015).

46. Stephenson, J., Nutma, E., van der Valk, P. & Amor, S. Inflammation in CNS neurodegenerative diseases. Immunology 154, 204–219 (2018).

47. Doorn, K.J., et al. Microglial phenotypes and toll-like receptor 2 in the substantia nigra and hippocampus of incidental Lewy body disease cases and Parkinson’s disease patients. Acta neuropathologica communications 2, 90 (2014).

48. Ghadery, C., et al. Microglial activation in Parkinson’s disease using [(18)F]-FEPPA. J Neuroinflammation 14, 8 (2017).

49. Koshimori, Y., et al. Imaging Striatal Microglial Activation in Patients with Parkinson’s Disease. PLoS One 10, e0138721 (2015).

50. Varnäs, K., et al. PET imaging of [11C] PBR28 in Parkinson’s disease patients does not indicate increased binding to TSPO despite reduced dopamine transporter binding. European journal of nuclear medicine and molecular imaging 46, 367–375 (2019).

51. Mathys, H., et al. Author Correction: Single-cell transcriptomic analysis of Alzheimer’s disease. Nature 571, E1 (2019).

52. Grubman, A., et al. A single-cell atlas of entorhinal cortex from individuals with Alzheimer’s disease reveals cell-type-specific gene expression regulation. Nature neuroscience 22, 2087–2097 (2019).

53. Zhou, Y., et al. Human and mouse single-nucleus transcriptomics reveal TREM2-dependent and TREM2-independent cellular responses in Alzheimer’s disease. Nature medicine 26, 131–142 (2020).

54. Smith, A.M., et al. Diverse human astrocyte and microglial transcriptional responses to Alzheimer’s pathology. Acta Neuropathol 143, 75–91 (2022).

55. Benjamini, Y. & Hochberg, Y. Controlling the False Discovery Rate - a Practical and Powerful Approach to Multiple Testing. J R Stat Soc B 57, 289–300 (1995).

56. Felton, J.M., et al. Epigenetic Analysis of the Chromatin Landscape Identifies a Repertoire of Murine Eosinophil-Specific PU.1-Bound Enhancers. J Immunol 207, 1044–1054 (2021).

57. Langmead, B., Trapnell, C., Pop, M. & Salzberg, S.L. Ultrafast and memory-efficient alignment of short DNA sequences to the human genome. Genome Biol 10, R25 (2009).

58. Heinz, S., et al. Simple combinations of lineage-determining transcription factors prime cis-regulatory elements required for macrophage and B cell identities. Mol Cell 38, 576–589 (2010).

59. Kent, W.J., et al. The human genome browser at UCSC. Genome Res 12, 996–1006 (2002).

60. Notredame, C., Higgins, D.G. & Heringa, J. T-Coffee: A novel method for fast and accurate multiple sequence alignment. J Mol Biol 302, 205–217 (2000).

61. Erb, I., et al. Use of ChIP-Seq data for the design of a multiple promoter-alignment method. Nucleic Acids Res 40, e52 (2012).

62. Waterhouse, A.M., Procter, J.B., Martin, D.M., Clamp, M. & Barton, G.J. Jalview Version 2--a multiple sequence alignment editor and analysis workbench. Bioinformatics 25, 1189–1191 (2009).

63. Tamura, K., Stecher, G. & Kumar, S. MEGA11: Molecular Evolutionary Genetics Analysis Version 11. Mol Biol Evol 38, 3022–3027 (2021).

64. Bailey, T.L., Johnson, J., Grant, C.E. & Noble, W.S. The MEME Suite. Nucleic Acids Res **43**, W39–49 (2015).

65. Bailey, T.L. & Grant, C.E. SEA: Simple Enrichment Analysis of motifs. bioRxiv (2021).

66. Love, M.I., Huber, W. & Anders, S. Moderated estimation of fold change and dispersion for RNA-seq data with DESeq2. Genome Biol 15, 550 (2014).

67. Duan, L., et al. PDGFRbeta Cells Rapidly Relay Inflammatory Signal from the Circulatory System to Neurons via Chemokine CCL2. Neuron 100, 183–200 e188 (2018).

68. Stuart, T., et al. Comprehensive Integration of Single-Cell Data. Cell 177, 1888–1902 e1821 (2019).

69. Khozoie, C., et al. scFlow: A Scalable and Reproducible Analysis Pipeline for Single-Cell RNA Sequencing Data. (Authorea Preprints, 2021).

70. Tsartsalis, S., et al. Single nuclear transcriptional signatures of dysfunctional brain vascular homeostasis in Alzheimer’s disease. (2021).

71. Ewels, P.A., et al. The nf-core framework for community-curated bioinformatics pipelines. Nat Biotechnol 38, 276–278 (2020).

72. Langfelder, P. & Horvath, S. WGCNA: an R package for weighted correlation network analysis. BMC Bioinformatics 9, 559 (2008).

73. Zhang, B., Kirov, S. & Snoddy, J. WebGestalt: an integrated system for exploring gene sets in various biological contexts. Nucleic Acids Res 33, W741–748 (2005).

74. Al-Izki, S., et al. Practical guide to the induction of relapsing progressive experimental autoimmune encephalomyelitis in the Biozzi ABH mouse. Mult Scler Relat Disord 1, 29–38 (2012).

75. Baker, D. & Amor, S. Publication guidelines for refereeing and reporting on animal use in experimental autoimmune encephalomyelitis. J Neuroimmunol 242, 78–83 (2012).

76. Maggi, P., Sati, P. & Massacesi, L. Magnetic resonance imaging of experimental autoimmune encephalomyelitis in the common marmoset. J Neuroimmunol 304, 86–92 (2017).

77. Nardo, G., et al. Transcriptomic indices of fast and slow disease progression in two mouse models of amyotrophic lateral sclerosis. Brain : a journal of neurology 136, 3305–3332 (2013).

78. Nardo, G., et al. Immune response in peripheral axons delays disease progression in SOD1(G93A) mice. J Neuroinflammation 13, 261 (2016).

79. Hampton, D.W., et al. HspB5 Activates a Neuroprotective Glial Cell Response in Experimental Tauopathy. Frontiers in neuroscience 14, 574 (2020).

80. Hampton, D.W., et al. Cell-mediated neuroprotection in a mouse model of human tauopathy. J Neurosci 30, 9973–9983 (2010).

81. Torvell, M., et al. A single systemic inflammatory insult causes acute motor deficits and accelerates disease progression in a mouse model of human tauopathy. Alzheimers Dement (N Y) 5, 579–591 (2019).

82. Haenseler, W., et al. Excess alpha-synuclein compromises phagocytosis in iPSC-derived macrophages. Sci Rep 7, 9003 (2017).

83. Haenseler, W., et al. A Highly Efficient Human Pluripotent Stem Cell Microglia Model Displays a Neuronal-Co-culture-Specific Expression Profile and Inflammatory Response. Stem Cell Reports 8, 1727–1742 (2017).

